# Mechanism of DNA surface exploration and operator bypassing during target search

**DOI:** 10.1101/2020.01.29.924647

**Authors:** E. Marklund, B. van Oosten, G. Mao, E. Amselem, K. Kipper, A. Sabantsev, A. Emmerich, D. Globisch, X. Zheng, L. C. Lehmann, O. Berg, M. Johansson, J. Elf, S. Deindl

## Abstract

Many proteins that bind specific DNA sequences search the genome by combining three dimensional (3D) diffusion in the cytoplasm with one dimensional (1D) sliding on non-specific DNA^1–5^. Here we combine resonance energy transfer and fluorescence correlation measurements to characterize how individual *lac* repressor (LacI) molecules explore DNA during the 1D phase of target search. To track the rotation of sliding LacI molecules on the microsecond time scale during DNA surface search, we use real-time single-molecule confocal laser tracking combined with fluorescence correlation spectroscopy (SMCT-FCS). The fluorescence signal fluctuations are accurately described by rotation-coupled sliding, where LacI traverses ~40 base pairs (bp) per revolution. This distance substantially exceeds the 10.5-bp helical pitch of DNA, suggesting that the sliding protein frequently hops out of the DNA groove, which would result in frequent bypassing of target sequences. Indeed, we directly observe such bypassing by single-molecule fluorescence resonance energy transfer (smFRET). A combined analysis of the smFRET and SMCT-FCS data shows that LacI at most hops one to two grooves (10-20 bp) every 250 μs. Overall, our data suggest a speed-accuracy trade-off during sliding; the weak nature of non-specific protein-DNA interactions underlies operator bypassing but also facilitates rapid sliding. We anticipate that our SMCT-FCS method to monitor rotational diffusion on the microsecond time scale while tracking individual molecules with millisecond time resolution will be applicable to the real-time investigation of many other biological interactions and effectively extends the accessible time regime by two orders of magnitude.

---

Sequence-specific binding and recognition of DNA target sites by proteins such as transcription factors, polymerases, and DNA-modifying enzymes is at the core of cellular information processing. However, the ‘target search problem’ of how to rapidly yet accurately find a specific target sequence remains incompletely understood. One aspect of the search problem is addressed by the facilitated diffusion theory^1^, which postulates that proteins search the genome by combining 3D diffusion in the cytoplasm with 1D sliding on non-specific regions of the DNA^1,5^, effectively expanding the target region to the sliding distance. While facilitated diffusion has been thoroughly explored both theoretically and experimentally^2,6^, very little is known about the mechanism of how proteins explore the DNA surface during sliding. In order to explain the differences in association rate observed *in vivo* for binding to different operators, DNA binding proteins have been suggested to be able to bypass operators during sliding^2^. However, such operator bypassing has never been observed directly. It is also not known if the sliding protein redundantly samples each base, as would be expected for faithful 1D diffusion, or if the protein trades redundancy for speed and occasionally skips bases. In this study, we shed light on the important trade-off between sliding speed and accuracy in target site recognition by measuring how a prototypical DNA-binding protein, the transcription factor LacI, explores the DNA surface during sliding.

As a first step towards the detection of operator bypassing, we first established a single-molecule fluorescence resonance energy transfer (FRET)^7,8^ assay to directly monitor the kinetics with which LacI binds to its natural *lacO*_1_ operator site (*O*_1_). We generated a DNA construct containing *lacO*_1_ with a Cy5 FRET acceptor dye 5 bp from the *lacO*_1_ edge. The DNA was surface-immobilized, and individual DNA molecules were monitored with a total-internal-reflection fluorescence (TIRF) microscope (Fig. 1a). Upon addition of LacI labelled with rhodamine in the DNA-binding domain (LacI-R, Extended Data Table 1, Extended Data Fig. 1), its specific binding (Extended Data Fig. 2) to *lacO*_1_ led to a sudden appearance of fluorescence signals and FRET (Fig. 1b). LacI containing a single donor (one-step photobleaching) on the distal or proximal monomer subunit gave rise to a FRET distribution for binding events with peaks at FRET = 0.16 ± 0.001 and 0.89 ± 0.002, respectively (Fig. 1c and Methods). The rate of operator binding (*k*_on,obs_) depended on both LacI and Na^+^ concentrations (Fig. 1d and Extended Data Fig. 2g) and the rate of LacI dissociation (*k*_off,obs_) increased with increasing Na^+^ concentration (Extended Data Fig. 2d,g). Importantly, LacI-R retained the ability to bind to both naturally occurring *lacO*_1_ and *lacO*_3_ operators with binding affinities essentially identical to the ones previously reported for the native protein^9^ (Fig. 1e), indicating that dye labelling of LacI-R did not substantially impact on its operator binding. The DNA binding affinity of LacI-R was also compared to a construct where LacI was labelled further away from the DNA binding domain (LacI-Far). The two labelling variants displayed identical sliding speeds (Extended Data Fig. 1c,d) and very similar affinities for binding to *lacO*_1_ (0.5 and 1 nM for LacI-Far and LacI-R respectively, Extended Data Fig. 1e), further showing that the label impairs neither the specific nor the non-specific DNA binding of the protein. Interestingly, FRET time traces of LacI binding events exhibited instantaneous transitions between the proximal and distal binding orientations on the same operator (Fig. 1f). These ‘flipping’ transitions presumably arise from microscopic dissociation events, where initially operator-bound LacI slides away from the operator and undergoes a spontaneous flipping transition before rebinding the operator in a flipped orientation.

**Figure 1.**
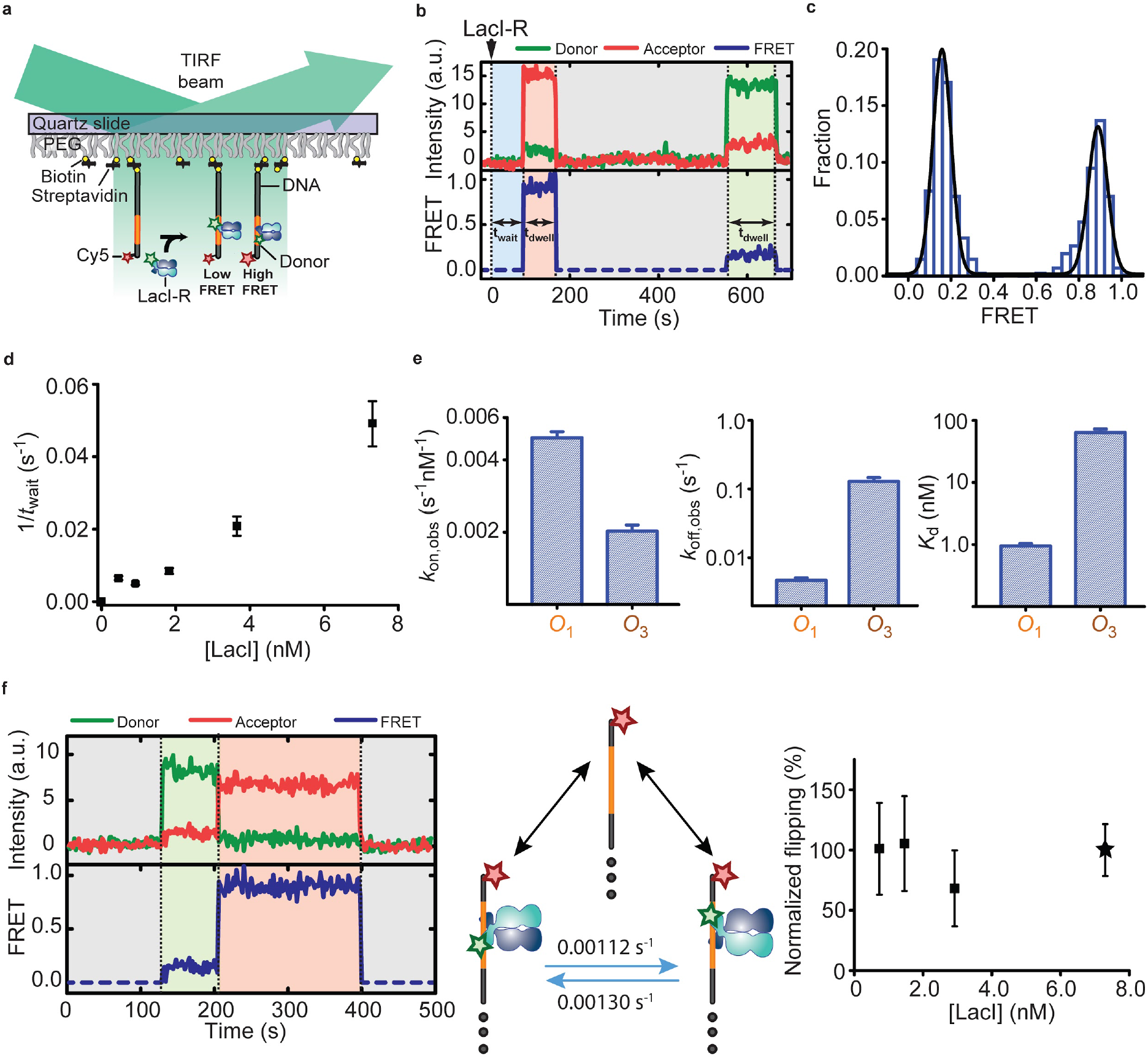
Monitoring LacI-operator interaction dynamics by single-molecule FRET. **a**, Schematic of FRET detection for specific LacI binding to its operator site (orange). The LacI dimer is labelled with the FRET donor, rhodamine (green star) and the DNA (grey) with the acceptor, Cy5 (red star). **b-e**, Analysis of specific binding to the *O*_1_ operator site. **b**, Donor fluorescence (green), acceptor fluorescence (red), and FRET (blue) traces showing specific binding of LacI containing a single donor on the proximal (left event) or distal (right event) monomer subunit. LacI-R was added at time zero (dotted line). The durations of the waiting phase (shaded blue) and dwell times (shaded red and green for high and low FRET values, respectively) are denoted as *t*_wait_ and *t*_dwell_, respectively. a.u., arbitrary units. **c**, FRET histogram from 6,500 specific binding events. The two peaks centred at FRET = 0.16 and 0.89 (derived from Gaussian fit, black line) result from the proximal and distal BR-labelling configurations. **d,**Dependence of the mean *t*_wait_ value for binding on LacI ([NaCl] = 1 mM) **e**, Rates of observed binding to (left: *k*_on,obs_, obtained as *t*_wait_^−1^*[LacI]^−1^) and dissociation from (middle: *k*_off,obs_) *O*_1_ and *O*_3_ sites and the resulting dissociation constants *K*_d_ (right). [LacI-R] = 7.3 nM, [NaCl] = 1 mM. **f**, Flipping transitions. Left: Donor fluorescence (green), acceptor fluorescence (red), and FRET (blue) traces showing a flipping transition where LacI-R transitions from a low FRET state (shaded green) to a high FRET state (shaded red). Centre: Cartoon schematic of the 3 distinct LacI binding states (see Methods) observed with a construct featuring a single *O*_1_ site. The rates for flipping transitions are 0.0011 ± 0.0004 s^−1^ (distal to proximal labelling configuration) and 0.0013 ± 0.0004 s^−1^ (proximal to distal configuration), errors represent 95% confidence intervals with at least 28,000 s of observed time in each state. Right: Flipping rates observed at various LacI-R concentrations, normalized to the flipping rate observed at a LacI concentration of 7.3 nM (asterisk), which was used for the determination of binding kinetics from single-site constructs as shown in (e). A 10-fold decrease in LacI-R concentration showed no significant change in flipping rate, suggesting that flipping events are not caused by the binding of a second LacI molecule. Error bars for *t*_wait_, *k*_off,obs_, *k*_on,obs_ and *K*_d_ are mean ± s.e.m. derived from at least 100 individual events.

In order to directly test for operator bypassing, we generated DNA constructs with two outer *lacO*_1_ sites separated by 30 bp of DNA that was either random (*ran*) or contained a third operator site (*lacO*_1_ or *lacO*_3_) (Fig. 2b). To discriminate between binding to the one *versus* the other outer *lacO*_1_ site, we used Cy5 and Alexa750 as distinct acceptor dyes. In the presence of LacI-R, fluorescence traces from individual *lacO*_1_-*ran-lacO*_1_ molecules exhibited spontaneous ‘switching’ transitions due to LacI sliding from one outer operator site to the other (Fig. 2a). These switching transitions involved a single LacI dimer, based on the following two observations. First, switching transitions were marked by fluorescence signals appearing in the one acceptor channel and simultaneously, i.e. given the camera frame rate of - 5 Hz, disappearing in the other acceptor channel. In stark contrast to these rapid changes, the corresponding mean waiting time for a single binding event at the same LacI concentration (Fig. 1d) was substantially longer with *t*_wait_= 199 s (see also Methods). Second, at this concentration, the frequency of switching transitions did not depend on the LacI concentration (Extended Data Fig. 3). We reasoned that a third, intervening operator should capture LacI sliding away from one of the outer *lacO*_1_ sites, thereby sequester it outside the FRET range (see also Supplementary Information 3.1.2) of either acceptor, and abolish the observed switching. Indeed, an intervening *lacO*_3_ or strong *lacO*_1_ site reduced the rate of switching (by a factor of 1.35 or 3.71, respectively), yet did not completely abolish it (Fig. 2b). Taken together, these observations show that a single LacI dimer can bypass intervening sites and slide between the outer *O*_1_ sites.

**Figure 2.**
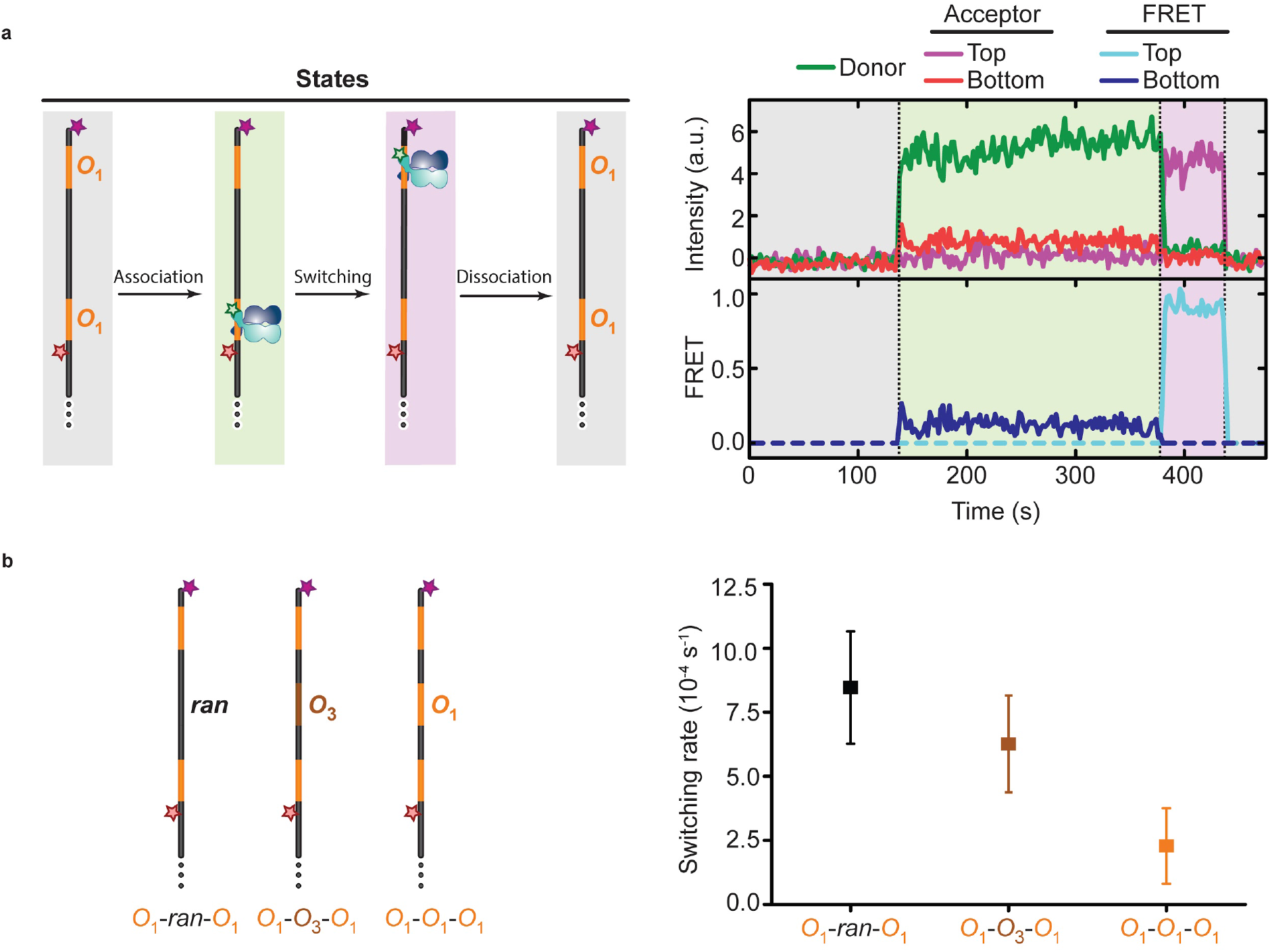
Direct observation of operator bypassing. **a**, Left: Schematic of DNA constructs with two outer *O*_1_ sites (orange) indicating the possible bound states of LacI-R. Top and bottom *O*_1_ sites are distinguished by Alexa750 (purple star) and Cy5 (red star) acceptor fluorophores, respectively. Right: Donor fluorescence (green), bottom (red) and top (purple) site acceptor fluorescence, as well as bottom (dark blue) and top site (light blue) FRET traces showing the transition of a single LacI dimer initially bound at the bottom site (shaded in green) switching to the top site (shaded in purple). In the absence of acceptor signals, the corresponding sections of FRET traces are set to zero, indicated by dashed lines for clarity. **b**, Left: Schematic of DNA constructs with two outer *O*_1_ sites (orange) and intervening random (*ran*) DNA (left panel) or an additional *O*_3_ (brown, middle panel) or *O*_1_ (right panel) site. Top and bottom *O*_1_ sites are distinguished by Alexa750 (purple star) and Cy5 (red star) acceptor fluorophores, respectively. Right: Quantification of the switching rates using HMM analysis^26^ (Extended Data Fig. 3) shown with 95% confidence intervals, derived from at least 40,000 seconds of observed binding time from at least 10 independent experiments.

To better understand the bypassing mechanism, we next sought to determine how LacI explores the DNA surface during target search. For this purpose, LacI-R was designed for bifunctional labelling to two proximal cysteines to reduce rotation of the dye relative to the protein. Indeed, peptide cleavage controls showed that essentially all LacI-R was homogeneously labelled with rhodamine dye bifunctionally attached to both adjacent cysteine residues (Extended Data Fig. 4a). We first characterized the orientation of individual labelled LacI-R molecules by measuring their polarization of fluorescence while sliding on flow-stretched *λ*-DNA (49 kB) using single-molecule wide-field epifluorescence and camera-based polarization detection^7,10–13^ (Fig. 3a, Extended Data Fig. 4b-j). These measurements showed a clear anisotropic polarization signal, implying a non-random fluorophore orientation during LacI sliding on DNA (Fig. 3a, Extended Data Fig. 4b-j, Supplementary Information 3.3). However, the limited temporal resolution (5 Hz) of these camera-based measurements could not resolve fast rotations of the protein around the DNA. Confocal fluorescence microscopy, on the other hand, has previously been used to study single-protein dynamics on a sub-millisecond time scale^14,15^. In order to more directly observe protein rotation around the DNA during sliding, we therefore combined real-time <u>s</u>ingle-<u>m</u>olecule <u>c</u>onfocal <u>l</u>aser <u>t</u>racking with <u>f</u>luorescence <u>c</u>orrelation <u>s</u>pectroscopy (SMCT-FCS) (Fig. 3b). This allowed us to monitor rotational diffusion on the microsecond time scale (Fig. 3b-f), at the same time as translational diffusion was tracked on the millisecond time scale. The translational movements, both parallel and perpendicular to the long axis of the DNA (Fig. 3d), of individual LacI-R molecules were tracked and used to classify them as sliders or non-sliders (i.e., protein stuck on the glass surface) (Methods and Supplementary Tables 1 and 2). For FCS analysis^16^, the photon emission was collected with nanosecond accuracy (Extended Data Fig. 5). We determined the autocorrelation function (ACF) of the fluorescence signal for molecules that bleached in a single step (Fig. 3e,f). If LacI sliding were coupled to its rotation around the DNA, the component of the ACF decay due to changes in the fluorophore orientation should be correlated with the rate of translational diffusion. A decrease in the translational diffusion rate is therefore expected to slow the decay of the autocorrelation in the relevant time regime. Indeed, when the experiment was repeated with a larger, maltose-binding protein fusion of LacI (LacI-MBP-R), we measured slower translational diffusion (0.027 ± 0.001 μm^2^s^−1^ *versus* 0.035 ± 0.002 μm^2^s^−1^, mean ± s.e.m; Fig. 3d, Extended Data Fig. 6a) as well as slower decay in the autocorrelation of the fluorescence signal in the 20 to 100 μs range (Fig 3e). We note that no such difference between LacI-R and LacI-MBP-R was observed when they were immobilized on the glass surface (Fig 3f, Extended Data Fig. 6b). The contributions to the autocorrelation decay due to dye photophysics or flexibility in rotational attachment were therefore essentially identical for the two proteins. We thus conclude that the difference in autocorrelation between small and large proteins was due to 1D diffusion. Our data therefore show that LacI sliding is coupled to its rotation around the DNA with characteristic decay times on the order of 40 μs.

**Figure 3.**
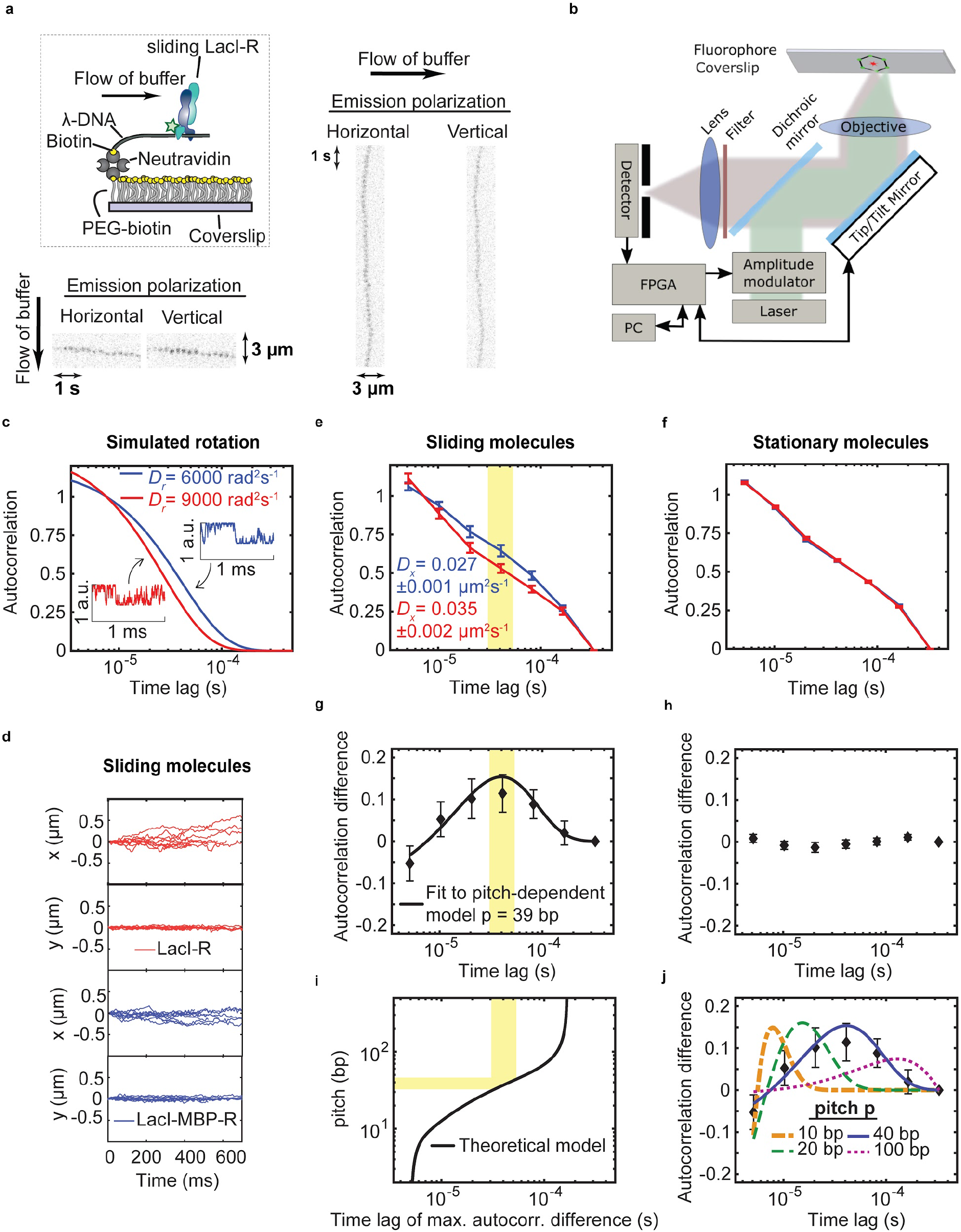
Pitch determination for rotation-coupled sliding on DNA. **a**, Schematic for the DNA flow-stretching experiment (top left). Kinetic series of images in horizontal and vertical emission polarization showing two representative sliding LacI molecules when DNA is stretched in the vertical (bottom left) or horizontal (right) direction. The DNA stretching direction is indicated in each example, showing vertical polarization emission when DNA is stretched in the vertical direction (bottom left) and flipped polarization emission when the stretching direction is flipped 90 degrees to be in the horizontal direction (right). **b**, Schematic of the SMCT-FCS setup. Black arrows: direction of communication between each block. Green dots: laser pattern during tracking of a fluorophore (red dot). **c**, Simulated traces (inset) of faster (red) and slower (blue) fluorophore rotation and their resulting autocorrelation functions. a.u., arbitrary units. **d**, x-(DNA direction) and y-coordinate of sliding LacI-MBP-R (blue, 151 molecules) and LacI-R (red, 91 molecules) 10 representative traces are shown for each species (see also Extended Data Fig. 6a). **e-f**, Mean normalized autocorrelation of the fluorescence signal for sliding (e) and stationary (f) LacI-MBP-R (blue) and LacI-R (red). Diffusion constants are averages from all sliding molecules. Error bars: standard errors. **g-h**, Difference in autocorrelation between LacI-MBP-R and LacI-R for sliding (g) and stationary (h) molecules. Error bars: standard errors. The black line in (g) shows the best fit to a rotation-coupled sliding model, while bootstrapping the trajectories yields an average pitch for the rotation of 39 ± 9 bp (mean ± s.e.m). **i**, Theoretical dependence of the pitch of rotation coupled sliding to the time lag of maximum difference in the autocorrelation functions. Area shaded in yellow: time regime corresponding to pitch values within the standard error of the experimentally measured pitch. **j**, Best fits of the rotation-coupled sliding model when the pitch is constrained at four different levels.

To estimate the bp distance that LacI translocates per revolution, we fit the difference in autocorrelation to a model where the pitch of the helical rotation is the only free parameter (Fig. 3g-j, Extended Data Fig 7). The fitting method accurately returned the correct pitches when it was tested on theoretical rotational autocorrelation functions, convoluted with the background noise processes obtained from the stationary molecules (Extended Data Fig. 7a,b). For the experimental data, fitting resulted in a pitch estimate of 39 ± 9 bp (Fig. 3g, see Extended Data Fig. 7c for individual repeats). To explore the signal-to-noise ratio in our SMCT-FCS experiments, we carried out simulations of fluorophore rotation using the experimentally estimated pitch as well as the same amount of data, shot noise, and data filtering steps as in the experiments. Notably, the resulting simulated differences in autocorrelation were very similar to the experimentally determined ones (Extended Data Fig. 7d and Methods), confirming that our SMCT-FCS experiments yielded signal amplitudes and associated errors well within the range of what could be expected from theory. We conclude that the sliding protein does not faithfully track the DNA helix but instead slides with a longer pitch. A sliding model where LacI sometimes slips between grooves *via* microscopic hops would agree with these observations and would also contribute to operator bypassing.

To determine which microscopic parameters for hopping, i.e., hop length and frequency, are consistent with the experimental observables, i.e., switching rate, flipping rate, and pitch, we simulated the processes for broad ranges of microscopic parameters (Fig. 4a,b). Flipping (Fig. 4c) and switching (Fig. 4d) rates were sampled in simulations corresponding to the FRET experiments (see Methods). Since both the overall 1D diffusion rate and the pitch of rotational sliding are known from the SMCT-FCS experiments, the hop frequency *k* can be calculated for each hop length *x* (see Fig. 4b). The absolute hop and switching rates also depend on how often LacI dissociates from the operator (Fig. 4c,d), since LacI cannot hop if it is bound to the operator site. The experimental flipping rate (dashed line, Fig. 4c) represents a lower bound for the absolute hop rate, since LacI cannot flip without hopping. At the same time, the absolute rate of the switching transitions defines a relation between the hop length and the operator dissociation rate, where longer hops that can bypass the intervening operator are compensated for by more frequent dissociation (dashed line, Fig. 4d). By determining where the dashed lines in Fig. 4c,d overlap (see Methods) to combine the different experimental constraints (Fig. 4e), we find that the average hop length cannot exceed 16 ± 8 bp, corresponding to a minimum hop frequency of 4 ± 1 ms^−1^ (S.E.M from bootstrapping). We note that our model does not make any assumptions or predictions about the rate for the microscopic association of LacI to its operator sites, since the overall results are insensitive to this parameter (Extended Data Fig. 8).

**Figure 4.**
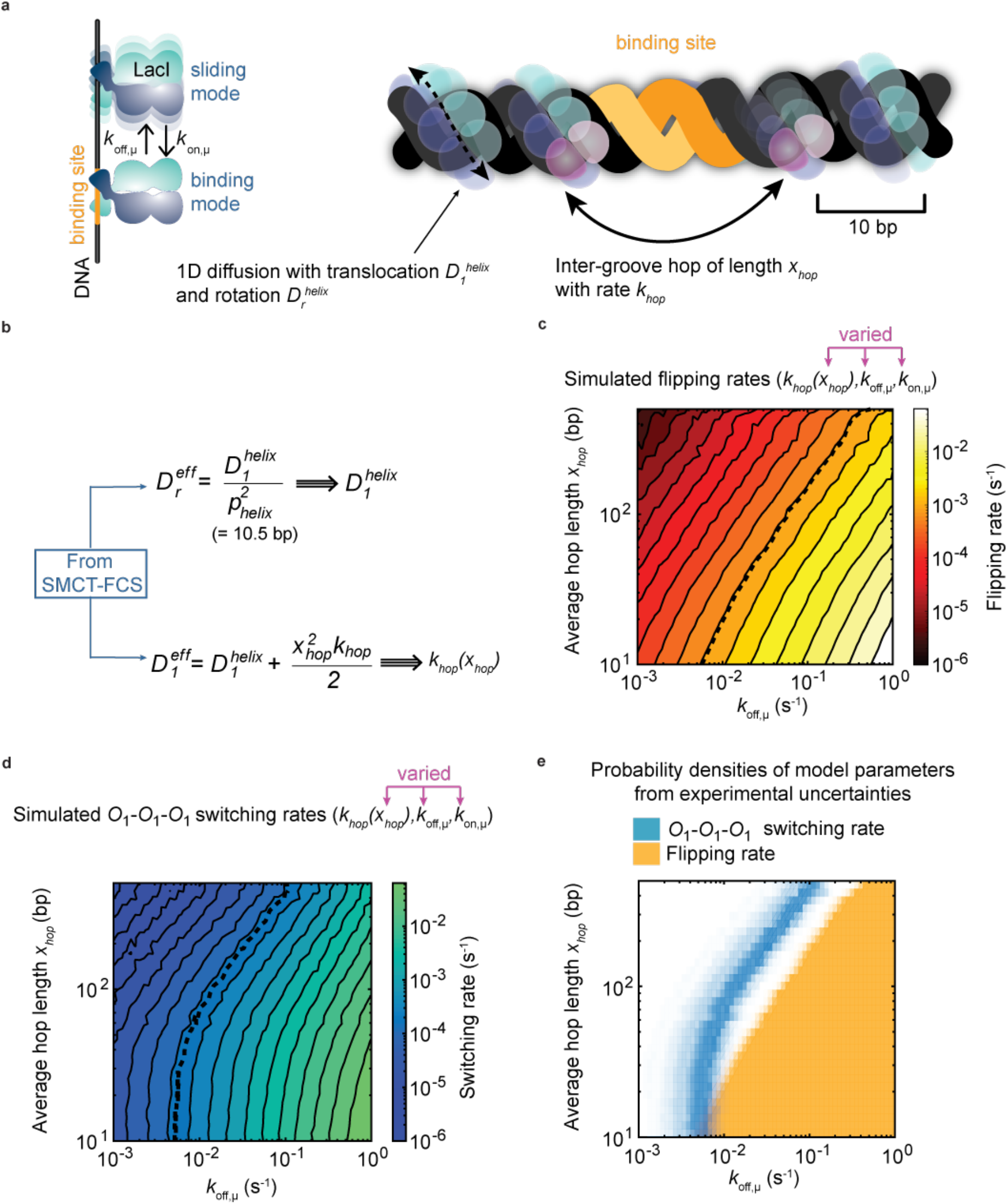
Determination of hop length and frequency. **a**, Cartoon schematic of target search mechanism where dimeric LacI (blue and cyan spheres) combines faithful DNA groove tracking (1D diffusion) with short and frequent inter-groove hops that can bypass a specific binding site (orange). **b**, Equations describing how the effective rotational (*D*_*r*_) and translational (*D*_*1*_) diffusion coefficient from SMCT-FCS are used to set parameters (*k_hop_(x_hop_)*) in the simulations of the FRET experiments. **c-d**, Flipping (c) and switching (d) rates calculated from stochastic simulations as a function of the model parameters. The dashed lines represent the experimentally observed rate of flipping on the same operator (c) and switching between two outer operators with an intervening operator in the middle (d) **e**, Model parameters compatible with the observed rates for switching (blue) and flipping (orange). The transparency values of the surfaces are scaled according to the probability density for the parameters given the experimental data. The flipping rate surface does not decrease with increasing values of *k*_*off,μ*_, since the simulations yield the maximum possible flipping rate. In other words, parameters corresponding to a higher maximum flipping rate could have also generated a lower actual flipping rate. The microscopic association rate from the sliding state to the operator (*k*_*on,μ*_) was infinite in these simulations, see Extended Data Fig. 8 for simulations with finite values of *k*_*on,μ*_.

Our measurements of protein-DNA interactions on the microsecond time scale show that the *lac* repressor rotates while sliding, and with a pitch that exceeds the 10.5-bp DNA pitch as a consequence of frequent and short hops. Such hopping may result from the non-specific binding that is sufficiently weak to optimize overall search speed, while at the same time inevitably leading to frequent operator bypassing. Operator bypassing does not necessarily reduce the total probability of rapidly binding the specific site, since 1D diffusion is redundant and involves many revisits to the same bases^1^. In fact, the observed hopping frequency allows for rapid scanning of the DNA and speeds up the first encounter with the specific site by ~100% in stochastic simulations of target search, despite the frequent bypassing events (Supplementary Information 4.5.3). Frequent and short hopping should also enable proteins to bypass obstacles encountered during sliding along the DNA. Indeed, such bypassing has been observed for proteins sliding on nucleic acids^17^. Although hopping has not been resolved in these cases, it may represent a common property harnessed by DNA-binding proteins to overcome obstacles that would block passage if sliding were strictly confined to rotation-coupled 1D diffusion.

Finally, we anticipate that our combination of confocal single-molecule tracking and FCS (SMCT-FCS) will lend itself to characterising many molecular interactions hundreds of times faster than what has been accessible hitherto^18^.

**Supplementary Information** is linked to the online version of the paper.

## Acknowledgements

We thank Paul Blainey and Kan Xiong for helpful advice on the flow-stretching assay, Chirlmin Joo for helpful discussions, Joakim Laksman for initial work on analysis code, and David Fange, Irmeli Barkefors and Måns Ehrenberg for helpful comments on the manuscript. The project was funded by the European Research Council (ERC), the Swedish Research Council (VR) and the Knut and Alice Wallenberg Foundation (KAW). All raw data and analysis software that were developed for this project are available from the corresponding authors (J.E. and S.D.) upon reasonable request.

## Author Contributions

E.M., B.v.O, M.G., and E.A. contributed equally to this manuscript. J.E. conceived of the confocal tracking with polarization readout concept and S.D. conceived of the smFRET approach to directly observe operator bypassing; J.E., E.M. and E.A. conceived of the SMCT-FCS implementation; S.D., B.v.O., M.G., J.E., and A.S. designed the smFRET study; E.A. developed and built the SMCT-FCS microscope; S.D., E.M., and K.K. developed the purification and labelling scheme. E.M. and L.C.L. purified and labelled LacI. E.M. implemented the fluidic assay, with input from M.J. E.M. carried out all tracking experiments, with assistance from E.A., K.K., and X.Z. E.M. and E.A. developed theoretical models and data analysis methods for SMCT-FCS tracking experiments. E.M. carried out analysis of SMCT-FCS tracking experiments. M.G. generated all smFRET DNA constructs and collected smFRET data with B.v.O. B.v.O. analysed FRET time traces with assistance from M.G. B.v.O. carried out HMM analyses of FRET time traces. A.E. provided initial code. D.G. supported HPLC purification and CNBr-mediate cleavage experiments. E.M. designed and carried out stochastic simulations, with input from J.E, O.B., and S.D. S.D., J.E., E.M., and E.A. wrote the paper, with input from all authors.

## Author Information

The authors declare no competing financial interests. Correspondence and requests for materials should be addressed to J.E. and S.D.

## Online Methods

### Protein purification, labelling and functionality testing

All LacI constructs (Extended Data Table 1) featured a C-terminal 6xHis-tag for affinity purification and a truncated tetramerisation domain, a design that previously has been shown to retain the ability to form dimers^19^. Additional cysteines were introduced into the amino acid sequence to enable bifunctional labelling. Detailed descriptions of the purification and labelling can be found in the Supplementary Information 1. All single-molecule measurements were carried out with the same batch of LacI-MBP-R and LacI-R. The affinity for LacI-R binding to the *O*_1_ and *O*_3_ operators was determined using single-molecule FRET measurements (Fig. 1e, Extended Data Fig. 1e).

### DNA constructs for smFRET

Double-stranded DNA constructs that contained operator sites as indicated in Extended Data Table 1, the FRET acceptors (Integrated DNA Technologies) Cy5 (backbone-incorporated) and/or Alexa750 (attached to position 5 of a dT base *via* a 6-carbon linker), and an end-positioned biotin moiety were generated by annealing and ligating a set of overlapping, complementary oligonucleotides. High-performance liquid chromatography (HPLC)-purified oligonucleotides were mixed at equimolar concentrations in 50 mM Tris pH 8.0, 100 mM KCl, 1 mM EDTA, annealed with a temperature ramp (95–3°C), ligated with T4 DNA ligase (New England Biolabs), and purified by PAGE. Successful ligation was confirmed by denaturing PAGE.

### DNA flow stretching

Preparation of flow channels as well as tethering and stretching of dsDNA by laminar flow was carried out according to previously published methods^20,21^. The 49 kB *λ*-DNA, anchored on one end to a poly[ethylene glycol] (PEG)-passivated coverslip^20^ *via* a biotin-neutravidin linkage, and a fluorescent protein concentration between 10 and 100 pM were used for sliding experiments (see Supplementary Information 2.1 for details).

### Single-molecule fluorescence resonance energy transfer (FRET) microscopy

Biotinylated and fluorophore-labelled DNA constructs were surface-immobilized on PEG-coated quartz microscope slides through biotin-streptavidin linkage^22,23^. Cy3 and Cy5 fluorophores were excited with 532 nm Nd:YAG and 638 nm diode lasers, respectively, and fluorescence emissions from Cy3, Cy5, and Alexa750 were detected using a custom-built prism-based TIRF microscope, filtered with ZET532NF (Chroma) and NF03-642E (Semrock) notch filters, spectrally separated by 635 nm (T635lpxr) and 760 nm (T760lpxr) dichroic mirrors (Chroma), and imaged onto the three thirds of an Andor iXon Ultra 888 EMCCD camera. Data acquisition was controlled using MicroManager. Fluorescence emission time traces were corrected to account for the direct excitation of Cy5 by the 532 nm laser as well as for the bleedthrough from the Cy3 into the Cy5 channel and that from Cy5 into the Alexa750 channel. Data were analysed using the Fiji distribution of ImageJ, IDL, and Matlab.

Imaging experiments were carried out in imaging buffer containing 100 mM 4:1 K_2_HPO_4_:KH_2_PO_4_ pH 7.4, 1 mM 2-Mercaptoethanol, 0.05 mM EDTA, 100 μg/ml acetylated BSA (Promega), 10% (v/v) glycerol, 10% (w/v) glucose, 0.01% Tween 20 (v/v), 2 mM Trolox to reduce photoblinking of the dyes^24^, an enzymatic oxygen scavenging system (composed of 800 μg/ml glucose oxidase and 50 μg/ml catalase), as well as 1-160 mM NaCl (as indicated). LacI was supplied by infusing the sample chamber with imaging buffer supplemented with 0.5-7.3 nM LacI (as indicated) using a syringe pump (Harvard Apparatus). For measurements of *t*_wait_, the duration of the waiting period after supplying LacI and prior to the first binding event, as shown in Fig. 1b, was recorded. For all other measurements, including the determination of dwell times and switching rates, data were recorded on multiple fields of view in the presence of 1 mM NaCl and 0.9 nM donor-labelled LacI.

### Single-molecule wide-field fluorescence polarization microscopy

Schematics of the optical setup are shown in Extended Data Fig. 4b and in more detail in Extended Data Fig. 5a. A supercontinuum laser (NKT EXW-4), spectrally filtered using a combination of Semrock FF01-770SP-25 and Semrock FF01-532-18 filters, was used for fluorophore excitation at 540 ± 18 nm. The light was brought to the custom built microscope through a multimode fibre (Thorlabs M69L05) with a mode scrambler. The mode scrambler both randomizes the polarization and diffuses the light source. The sample was excited through an EPI-illumination configuration using a dichroic mirror (Semrock FF552-Di02) and a 100x Nikon objective (CFI Plan Apo Lambda, NA = 1.45). Fluorescence was collected using a dichroic mirror (Semrock FF552-Di02) and filter (Semrock FF01-585-40), which was followed by a 2x expander (Thorlabs achromatic doublets f = 100 mm and f = 200 mm) to obtain a final total magnification of 200x and a pixel size of 80 nm. The emitted light was divided into its horizontal and vertical components by a polarized beam splitter in between two relay lenses (Thorlabs achromatic doublets f = 150 mm). The two polarization components were imaged onto the two halves of an EMCCD camera (Princeton Instruments PhotonMAX 512). The laser intensity at the sample was set to 3 kW/cm^2^ and the EMCCD was configured to have an exposure time of 160 ms, resulting in a 5 fps frame rate. The EMCCD gain was optimized to obtain an average signal-to-noise ratio of 1.6.

### Peptide cleavage assay with CNBr

The assay was performed according to a previously published protocol^25^. Labelled protein samples were incubated in 0.25 M CNBr for 24 h before being lyophilized, resolubilised, and analysed by SDS-PAGE.

### Single-molecule confocal laser tracking with fluorescence correlation spectroscopy (SMCT-FCS)

#### Optical setup

Schematics of the optical setup are shown in Fig. 3b of the main text and in more detail in Extended Data Fig. 5a. After spectral filtering, the laser light was subjected to an amplitude modulator consisting of two polarizers with a pockel cell (Newport 4102NF) in between. A spatial filter (11 mm lens, 10 μm pinhole followed by a 75 mm lens) was used to obtain a pure TEM00 mode. A tip tilt piezo mirror (Piezosystem Jena PSH 10/2) was placed after the dichroic mirror at the back focal plane of the objective. The excitation laser was set to circular polarization in the xy-plane by rotating a combination of a half and quarter wave plates and measuring the polarization after the objective. The intensity at the back focal plane of the objective was set to 15 μW, and emitted fluorescence was collected through the dichroic mirror followed by an imaging lens, a 75 μm pinhole, a polarizing beam splitter (PBS), and an avalanche photodiode (APD, SPCM-AQRH Excelitas) for each polarization. A field programmable gate array (FPGA, NI 7852R) was used for sampling each APD signal, time tagging photon arrival times, triggering laser excitation pulses, and controlling the real-time feed-back of the piezo tip/tilt mirror.

#### Single-molecule-tracking

The tracking system was started as a raster scan. The scan works both as a standard confocal imaging system and a search for potential tracking events. Here the excitation laser was triggered with a 500 μs pulse with a sampling rate of 1900 samples/s. After a short delay (520 ns) a detection window of 500 μs was opened, during which the FPGA was counting the photons from the APDs. The summed APD photon count within the window was then used for further analysis in the FPGA. If the photon count was higher than a set threshold, the system entered the tracking mode. In the tracking mode, the tip/tilt piezo mirror traced out a circle centred around the position where the photon count passed the threshold requirement with a revolution time of 4 ms and a diameter of 290 nm. During each 4 ms revolution, the laser was triggered six times during equally spaced intervals, each with a duration of 500 μs with a corresponding detection window. The fluorophore offset from current estimated positions was triangulated by calculating a rolling mean-weighted centroid using the 12 latest points. A PID controller used the offset value to generate a correction for the current position, followed by the triggering of a position update of the piezo tip/tilt mirror with the new, corrected fluorophore position. This was performed for each new measurement point, and the tracking scheme was repeated until the mean photon count dropped below a set threshold, terminating the tracking loop and bringing the system back into the raster scanning mode. Details on the optimization of tracking parameters can be found in the Supplementary Information 3.2.

#### Photon counting data acquisition

Our photon counting mechanism served two purposes. Firstly, it summed all events for a given time window for real-time adaptation of the tracking beam. Secondly, it time-tagged all photons with ns-accuracy and sent the data to the computer for storage and later FCS post-processing analysis. Each detection window of 500 μs had a 200 Mhz loop with counters. Every time a photon was detected by an APD, the system updated the counters; two counters were gradually increased and delivered to the centroid calculation loop the accumulated photon counts of each APD. Simultaneously, one counter kept track of the 200 Mhz loop and, when triggered, delivered the time stamp of that photon. This value together with the APD ID and a rollover flag was sent to the computer and saved. These time-tagged time-resolved (TTTR) data, which comprise of an asynchronous record of the photon times, could be used to reconstruct the photon time traces corresponding to tracking trajectories with a time resolution of 5 ns and were used for FCS analysis on individual single-molecule tracking trajectories.

#### SMCT-FCS system benchmark

To benchmark and test both the tracking and FCS part of the SMCT-FCS microscope, immobilized fluorescent beads where tracked while moving the stage in a circle, see Extended Data Fig. 5b and Supplementary Information 3.2.

### Data analysis

#### Analysis of single-molecule FRET time traces from single-site constructs

As per quantification of the number of photobleaching steps for LacI-R and LacI-MBP-R bound to O_1_, 69.4% of labelled LacI-R and 77.2% of labelled LacI-MBP-R dimers featured a single dye and exhibited one-step photobleaching; 30.6% of labelled LacI-R and 22.8% of labelled LacI-MBP-R featured two dyes, leading to two-step photobleaching. For simplicity, we focused our analyses on LacI containing a single donor dye (one-step photobleaching, see Supplementary Information 3.1 for details) on the distal or proximal monomer subunit to determine waiting and dwell times using DNA constructs with a single operator site. The resulting histogram of mean FRET values for binding events exhibited two distinct peaks centred at FRET = 0.16 and FRET = 0.89, respectively (Fig. 1c). To determine the rate of operator binding (*k*_on,obs_), we quantified the ‘waiting time’ (*t*_wait_ *= 1/{k*_on,obs_*\*[LacI]}*) after LacI addition before any detectable appearance of fluorescence signals. To measure dissociation rates of LacI from the operator (*k*_off,obs_), we measured the dwell times of binding events (*t*_dwell_=*1/k*_off,obs_) (Extended Data Fig. 2d,f). LacI dissociation events cannot be unambiguously distinguished from photobleaching. For this reason, we determined dwell times at multiple laser power densities to identify imaging conditions where photobleaching did not affect reported dwell times (Extended Data Fig. 2d,f).

#### Switching transitions observed in single-molecule FRET time traces

Switching transitions were monitored at a concentration of 0.9 nM donor-labelled LacI and involved a single LacI dimer only. The presence of more than one (labelled or unlabelled) LacI dimer during the observed switching transitions can be ruled out based on the following considerations. Switching transitions were marked by fluorescence signals rapidly appearing in the one and simultaneously disappearing in the other acceptor channel (see Fig. 2a and Supplementary Information 3.1.2 for details). In stark contrast to these rapid changes, the mean waiting time for a single binding event, measured at the same LacI concentration (Fig. 1d), was substantially longer with *t*_wait_= 199 s. From hidden Markov modeling (HMM) analysis, the probability of an unbinding event (p = 7.66 * 10^−3^) followed by another binding to the opposite flanking operator site (p = 1.51 * 10^−4^) is calculated as 1.15 * 10^−6^ s^−1^ for the *O*_1_*-O*_1_*-O*_1_ construct, less than 1% of the lowest observed switching rate. Moreover, the frequency of switching transitions did not depend on the LacI concentration (Extended Data Fig. 3). Thus, a single LacI dimer only underlies the observed switching transitions.

For the analysis of switching transitions, we included in our analyses LacI dimer with a single donor dye as well as LacI dimer with a donor on both LacI monomers. Based on Cy5 and Alexa750 fluorescence intensities as well as the corresponding FRET values, time traces recorded from individual *O*_1_*-ran-O*_1_, *O*_1_*-O*_1_*-O*_1_, or *O*_1_*-O*_3_*-O*_1_ molecules were first divided into segments corresponding to the following distinct states: 0: Unbound state, no LacI; 1: Bottom (Cy5) *O*_1_ site bound by a LacI dimer with a single donor dye distal to the Cy5 acceptor dye; 2: Bottom (Cy5) *O*_1_ site bound by a LacI dimer with a single donor dye proximal to the Cy5 acceptor dye; 3: Bottom (Cy5) *O*_1_ site bound by a LacI dimer with a donor on each of the two monomers; 4: Top (Alexa750) *O*_1_ site bound by a LacI dimer with a single donor dye distal to the Alexa750 acceptor dye; 5: Top (Alexa750) *O*_1_ site bound by a LacI dimer with a single donor dye proximal to the Alexa750 acceptor dye. 6: Top (Alexa750) *O*_1_ site bound by a LacI dimer with a donor on each of the two monomers.

#### Hidden Markov modelling (HMM) analyses

In order to quantify the frequency of switching transitions, we carried out hidden Markov modelling (HMM) analyses^26^. Time traces from individual 3-site DNA molecules, divided into segments corresponding to the distinct states as described above, were concatenated and analyzed using HMM with Gaussian emissions (hmmlearn package, Python) to determine FRET states and corresponding transition probabilities between each based on an optimizing Viterbi algorithm. Next, states 1-3 or 4-6 were grouped to assign LacI dimer as either bound to the bottom (Cy5) *O*_1_ site or the top (Alexa750) *O*_1_ site, respectively.

#### Analysis of wide-field fluorescence polarization microscopy data

Supplementary Information 3.3.

#### Trajectory building of confocal tracking data

The output data available for the real-time tracking part of the SMCT-FCS system is the tip/tilt mirror position for each 500 μs laser pulse, the total photon count during the same laser pulse interval and a flag showing if the tracking mode was active. In the parts of the datastream when the tracking mode was active, the weighted centroid position over 12 points (8 ms) was calculated to obtain x and y fluorophore positions for each time section. To avoid building trajectories of noise when no fluorophore was present in the confocal volume, a background threshold was calculated and used to remove trajectories when their signal was below the background. Sliding trajectories were classified using principal component analysis (PCA), where the classifier was optimized to effectively find sliders in flow channels with DNA but not in negative control channels without DNA (Extended Data Table 2). The eigenvalue of largest principal component of the trajectory position for sliders was at least 0.0015 μm^2^ and at most 0.0008 μm^2^ for stationary molecules (see Supplementary Table 2 for more classification parameters). Molecules bleaching in more than one discrete step were excluded from the data, so that only trajectories stemming from LacI dimers that were labelled with a single dye were included in the analysis. 1D diffusion constants for sliding trajectories were estimated using the covariance-based estimator^27^ from displacements (weighted centroid positions over 16 ms) along the principal component with the largest eigenvalue (the DNA stretching direction). Averages of diffusion constants are reported as 20% trimmed means with standard errors obtained by bootstrapping over the trajectories.

#### Autocorrelation functions from photon time tagging data

The second output available from the SMCT-FCS system is photon time tagging data, consisting of asynchronously recorded time-tagged time-resolved (TTTR) data points. Reconstructing a synchronous time trace of the photon time series would become very computationally and memory intensive and is therefore not feasible. Instead, we used a time-tag-to-correlation algorithm^28^ to directly convert the TTTR data to fluorescence correlation spectroscopy (FCS) curves

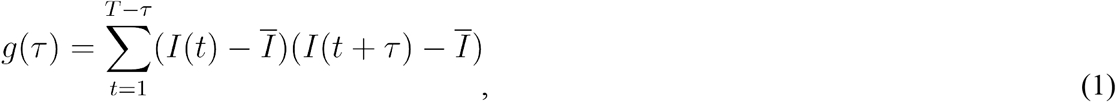

where *I(t)* is the photon count at time *t*, 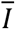 is the average photon count and *τ* is the time lag. This conversion was carried out with the combined TTTR data time series from both APDs, effectively calculating the FCS curves as if there were only one detector. The autocorrelation data are compensated for the 166 μs off time between laser pulses, i.e. we only used data during each 500 μs pulse. The normalized autocorrelation for each trajectory was calculated from the non-normalized autocorrelation *g(τ)* according to

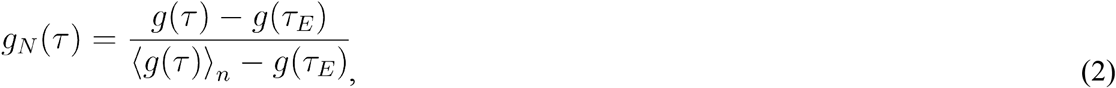

where *τ*_*E*_ is the last time lag of time scale and the < >_*n*_ operator denotes the average for the first *n* points of the time scale. For each sliding molecule the normalized autocorrelation function was calculated for seven time lags between 5 and 328 μs with *n* = 2 used for normalization. The average normalized autocorrelation for each protein and time point was then calculated as the mean of the single-molecule data. In the calculation of the mean, two criteria were used to avoid bias in the reported autocorrelation functions introduced by noisy and uninformative data. Firstly, autocorrelation functions for individual sliding molecules with a standard deviation of normalized autocorrelation for the different time lags larger than were excluded (which effectively discards autocorrelation functions that deviate heavily from the domain [0,1]). Secondly, autocorrelations stemming from molecules with a photon count rate lower than 5625 counts s^−1^ were also excluded.

#### Vector-based dipole point spread function model

The model used for the polarization and orientation dependent emission of a dipole is based on previous work^29^. Briefly, the polarization intensity in the back focal plane is found by translating the electric far field of the dipole into the back focal plane, and by integrating the intensity (square of the electric field) over the numerical aperture *NA* of the objective. After integration we obtain the fluorescence intensity

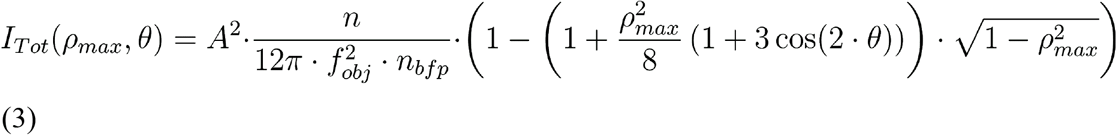

where *θ* is the polar angle of the dipole orientation, *A* is the amplitude of the dipole moment (dependent on the polarization of the excitation light and the dipole orientation), *n* and *n*_*bfp*_ are refractive indices surrounding the dipole and the back focal plane, respectively, *f*_*obj*_ is the focal length of the objective, *ρ*_*max*_ =*NA/n*, and *H(ρ_max_)* and *V*(*ρ*_*max*_) are diagonal matrices only dependent on *ρ*_*max*_ (see Supplementary Information 4.1 for derivation and details). The amplitude of the dipole moment was set according to the excitation light of the modelled process (*A* = *sin(θ)* for excitation light circularly polarized in the xy-plane, as in the SMCT-FCS measurements).

#### Simulating fluorescence traces and their autocorrelation functions

The relationship between the autocorrelation function *g(τ)* and the rotational diffusion constant *D*_*r*_ for rotation-coupled sliding was found by simulating fluorescence intensity traces emitted from dipoles rotating diffusionally around the DNA axis. Brownian dynamics simulations of the fluorophore rotation were performed for both 1D (see Supplementary 4.3) and 3D^30^ rotational diffusion, while the fluorescence intensity was calculated from the vector-based point-spread function model (Equation 3) at every time step. After calculating the autocorrelation function from intensity traces, we found that the simulated autocorrelations for 1D rotational diffusion could be described accurately with the model (Extended Data Fig. 7e)

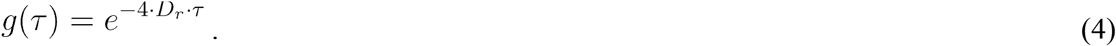

The simulation scheme was validated using the 3D rotational diffusion simulation, where the resulting autocorrelation, as expected, could be described accurately by an exponential with exponent −6*D_r_τ* (Extended Data Fig. 7e). Simulations of fluorophore rotation were also performed with shot noise to replicate the experimental conditions and to obtain a theoretical estimate for the error in our experimental signal (Extended Data Fig. 7d). To this end, the amount of emitted photons were sampled from a Poisson distribution at every time step, with average photon count rate, number of traces, and length of traces taken as the averages from the experiments.

#### Model fitting and pitch estimation

In rotation-coupled sliding, translational movement of the particle is coupled to rotation around the DNA. The rotational diffusion constant *D*_*r*_ and translational diffusion constant *D* are then related to each other according to

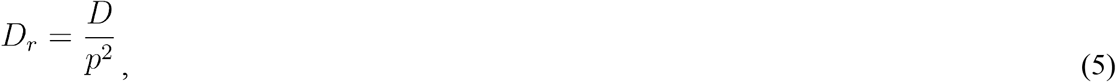

where *p* is the pitch of the rotational sliding, i.e. how far the particle translocates per angle it rotates around the DNA. To minimize contributions from the background (all contributions to the autocorrelation function that are independent of rotational sliding of the molecule, such as triplet state blinking or photons emitted by other light sources) and any hypothetical contribution from DNA rotation, model fitting was carried out using the difference in autocorrelation between two constructs of LacI (LacI-MBP-R and LacI-R) that were labelled with the same fluorophore but were sliding with different rates. The difference in autocorrelation was fitted to the following expression, with the normalization of the autocorrelation matching that applied to the experimental data,

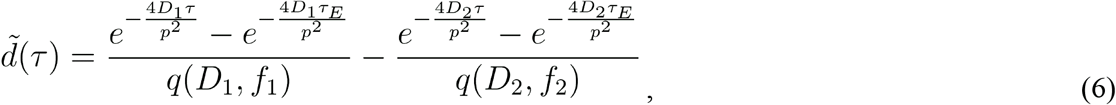

where the normalization factor *q* is defined as

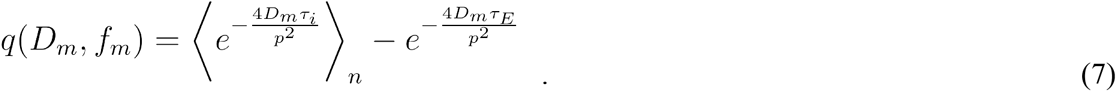

The free parameter of the model is the pitch *p*, which is fitted by minimizing the deviation between the experimental difference in autocorrelation and the theoretical difference according to equation 6 (see Supplementary Information 4.4 for details). Translational diffusion constants *D*_1_ and *D*_2_ were taken as the experimentally measured averages from the tracking data (20% trimmed means of the trajectory diffusion coefficients) but were allowed to vary in the fitting within their 95% confidence intervals from the experiment. In Fig. 3g and Extended Data Fig. 7c,d, the experimental data are plotted with the theoretical values according to equation 6. Fig. 3i depicts *p* as a function of the time lag of maximum autocorrelation difference. To obtain this theoretical curve, equation 6 was maximized with respect to *τ*, and for this implicit relationship *p* was plotted against *τ*. Rotational pitches were converted into the unit of bp per full revolution around the DNA in the main text.

#### Stochastic simulations of LacI flipping and operator switching

In the operator switching model, LacI has two binding modes when in contact with DNA: operator bound and non-specifically bound. In the non-specifically bound mode, LacI can slide helically with diffusion rate constant 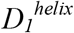, and undergo micro-hops of average length *x*_hop_ (sampled from a normal distribution with standard deviation *x*_hop_) with a rate of *k*_hop_. To obtain the maximum rate of flipping, we took into account that LacI has no memory of the initial flip orientation after a hop, meaning that LacI flips with a probability of 50% at every hop. When LacI has reached an operator, it can associate specifically to the operator with rate *k*_on,μ_. In the specific mode LacI can dissociate microscopically from the operator with rate *k*_off,μ_ to enter the non-specific mode. In both binding modes, LacI can dissociate macroscopically from the DNA with rate *k*_off,macro._ The effective diffusion constants for 1D translocation along the DNA and rotation around it are both sums of the diffusion constants for helically sliding and hopping respectively, and since LacI only rotates while sliding

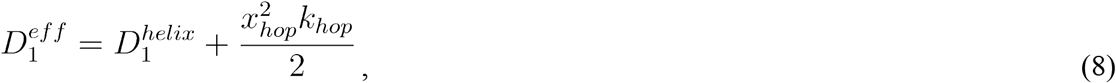

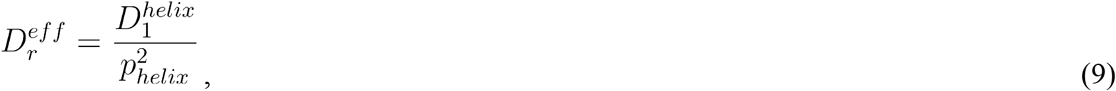

with 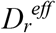, and 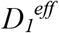 determined from the SMCT-FCS experiments, where the pitch of helical rotation *p*_*helix*_ is 10.5 bp per turn, and *x*_hop_ and *k*_hop_ are the only unknowns in equations 8 and 9, meaning that the hop rate is explicitly given by the average hop length. Switching and flipping events were simulated with Brownian dynamics on a discretized DNA (see Supplementary Information 4.5 for pseudo code) with reflective boundary conditions at the DNA edges. To achieve the same average trace length as in the experiments, *k*_off,macro_ was set to 0.008 s^−1^ for every set of the unknown parameters *Θ* = (*k*_on,μ_, *k*_off,μ_,*x*_hop_). This yielded the switching rate *s* and maximum flipping rate *f* as a function of *Θ*. To determine what set of parameters could generate the experimental data, the simulated and experimental switching and flipping rates were compared. This was done in a probabilistic manner to obtain a confidence interval associated with the experimental estimate of *x*_hop_, generating the probability densities seen in Fig. 4e. and Extended Data Fig. 8. For the switching rate *s* given the experimental traces *y,* that probability density is obtained from the relationship

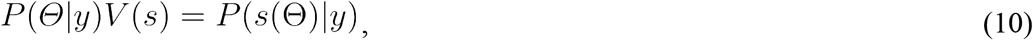

where the probability density *P*(*s*|*y*) is calculated by bootstrapping over the experimental traces. *V(s)* is the volume element in *Θ* space associated with each *s*, calculated from the *s(Θ)* obtained from simulations. With the simulated flipping rate (*f*) and the rate for switching on *O*_1_*-ran-O*_1_ DNA (*s*_*2*_) and *O*_1_*-O*_1_*-O*_1_ (*s*_*3*_), the joint probability distribution can be calculated as

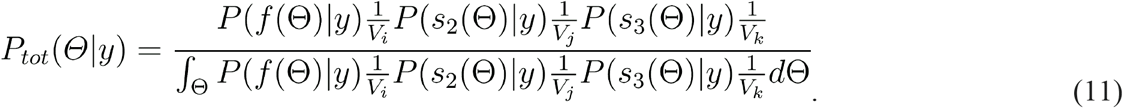

In practice, parameter estimation was carried out with *Θ* = (*k*_off,μ_,*x*_hop_), and the mean and standard error of *x*_hop_ were estimated for each *k*_on,μ_ in Fig. 4e and Extended Data Fig. 8. by marginalizing the joint distribution over *k*_off,μ_ and then calculating the mean and standard deviation of the marginalized distribution. The mean is

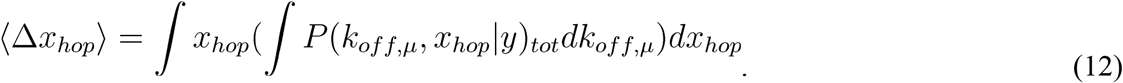

## Code availability

All software and analysis code developed for the project will be transferred upon reasonable request.

