## Extended Data for "Mechanism of DNA surface exploration and operator bypassing during target search"

### Extended Data Content:

Extended Data Figure 1: Analysis of LacI labelling and binding
Extended Data Figure 2: LacI binding kinetics
Extended Data Figure 3: Predominant switching and flipping transitions
Extended Data Figure 4: Camera-based polarisation measurements and characterisation of dye labelling
Extended Data Figure 5: Optical layout and calibration data for SMCT-FCS
Extended Data Figure 6: Traces captured with confocal tracking
Extended Data Figure 7: Fit of pitch-dependent autocorrelation model and repeats of SMCT-FCS experiments
Extended Data Figure 8: Probability densities of model parameters when taking into account the experimental FRET results
Extended Data Table 1: Sequences of LacI and single-molecule FRET DNA constructs
Extended Data Table 2: Sliding events in EMMCD and confocal tracking experiments

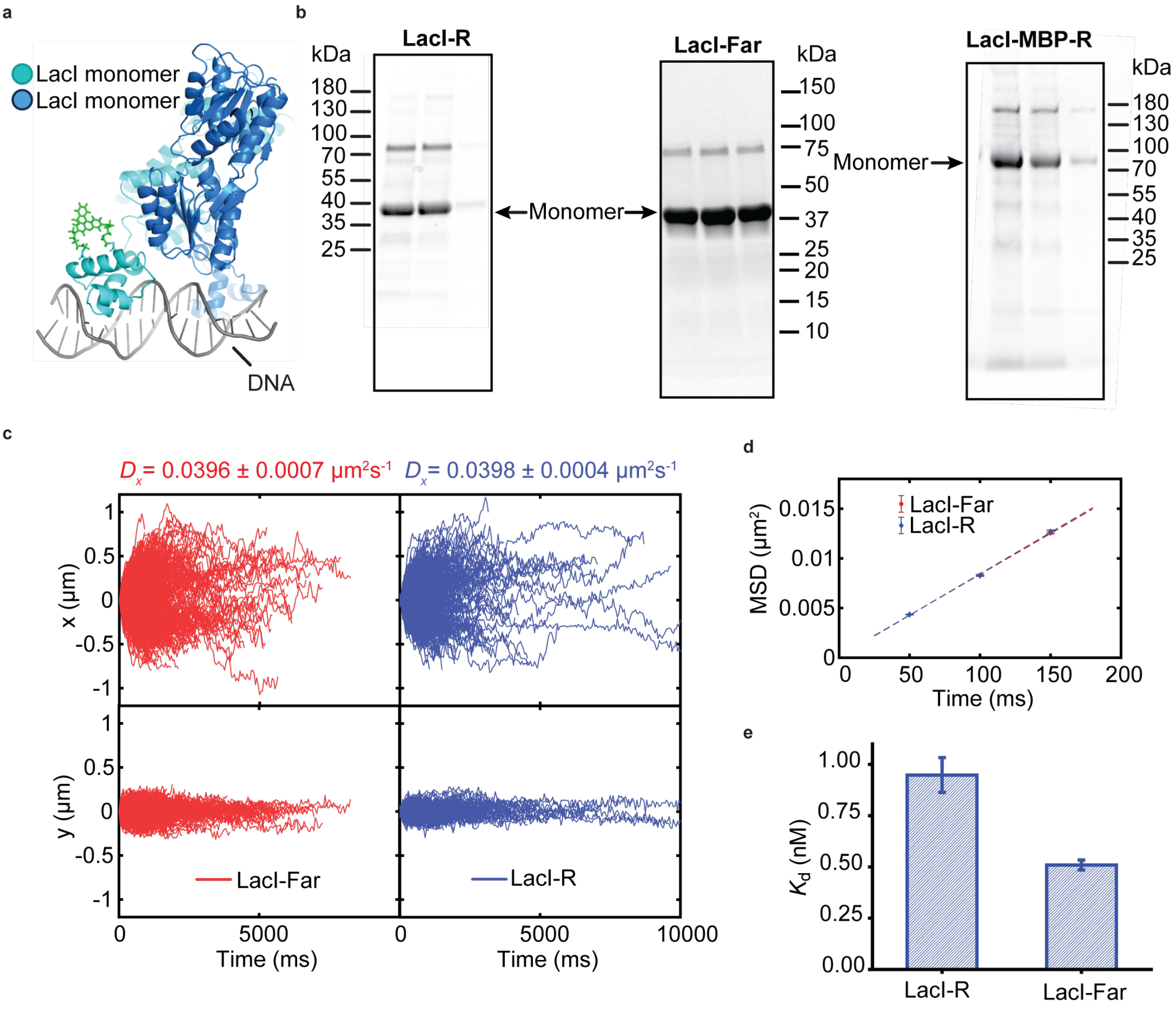


**Extended Data Figure 1: Analysis of LacI labelling and binding.** (**a**) Structural model (based on PDB code: 1OSL). LacI: blue/cyan, rhodamine: green, DNA: grey. (**b**) SDS-PAGE of labelled LacI fractions after dye removal visualized using rhodamine fluorescence. The bands corresponding to the monomeric sizes expected for LacI-R (left), LacI-Far (middle), and LacI-MBP-R (right) are indicated with blue arrows. The intensity of the monomeric band relative to the sum of the intensities of monomeric and dimeric bands is 77% or 86% for LacI-R or LacI-MBP-R, respectively. (**c**) x (DNA direction) and y coordinate of sliding LacI-Far (red) and LacI-R (blue) molecules obtained by EMCCD tracking with a 50 ms frame rate. In total, 779 and 409 sliding molecules were captured for LacI-Far and LacI-R, respectively. Diffusion constants along the x-coordinate are indicated as mean ± SEM. (**d**) Mean squared displacement for different time steps for LacI-R (blue) and LacI-Far (red). (**e**) The resulting dissociation constants $K_d$ for LacI-R and LacI-Far. [LacI] = 7.3 nM, [NaCl] = 1 mM.

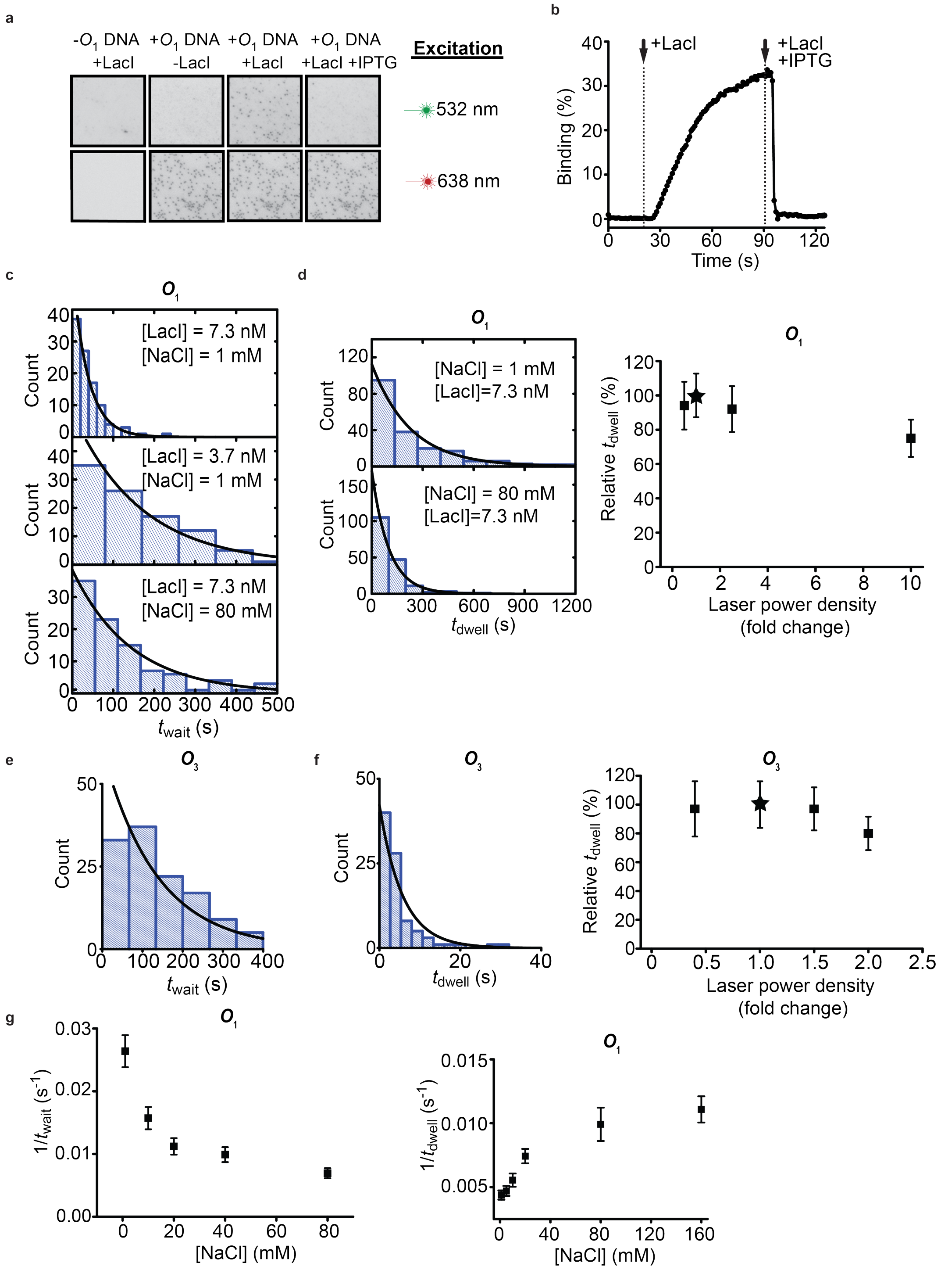


**Extended Data Figure 2. LacI binding kinetics.** (**a**) EMCCD images of acceptor channel under 532 nm (top: donor excitation) and 638 nm (bottom: acceptor excitation) illumination. Specific binding of individual donor-labelled LacI-R molecules (individual spots, top row) occurs in the presence (+$O_1$ DNA; individual spots, bottom row) but not in the absence (-$O_1$ DNA) of acceptor-labelled DNA containing an operator site. Addition of IPTG (+ IPTG) to the same field of view displaces specifically bound LacI. (**b**) Time trace of LacI-R occupancy in one field of view (> 1000 individual $O_1$ DNA molecules). 7.3 nM LacI-R were supplied after 20 s (first dotted line), followed by the addition of 7.3 nM LacI-R together with 100 mM IPTG after 90 s. (**c**) Histogram of $t_{wait}$ values for specific binding to $O_1$ ($N$ = 100 events) at various

LacI and NaCl concentrations as indicated. (**d**) Left: Histogram of $t_{dwell}$ values for specific binding to $O_1$ ($N$ > 100 events) at [LacI-R] = 7.3 nM and [NaCl] = 1 mM (top) or 80 mM (bottom). Right: Under standard imaging conditions (marked by the asterisk), measurements of $t_{dwell}$ for binding to $O_1$ are not affected by photobleaching. Shown is the mean relative $t_{dwell}$ for specific binding to $O_1$ ($N$ > 100 events) observed at [NaCl] = 1 mM and using various laser power densities. Dwell times and laser power densities were normalized to the standard imaging laser power density used for all other analyses (asterisk). (**e**) Histogram of $t_{wait}$ values for specific binding to $O_3$ ($N$ = 100 events) at [LacI] = 7.3 nM and [NaCl] = 1 mM. (**f**) Left panel: Histogram of $t_{dwell}$ values for specific binding to $O_3$ ($N$ = 100 events) at [LacI-R] = 7.3 nM and [NaCl] = 1 mM. Right panel: Under standard imaging conditions (marked by the asterisk), measurements of $t_{dwell}$ for binding to $O_3$ are not affected by photobleaching. Shown is the mean relative $t_{dwell}$ for specific binding to $O_1$ (N > 100 events) observed at [NaCl] = 1 mM and using various laser power densities. Dwell times and laser power densities were normalized to the standard imaging laser power density used for all other analyses (asterisk). (**g**) Dependence of the mean $t_{wait}$ (left) and $t_{dwell}$ (right) value for binding on NaCl ([LacI-R] = 7.3 nM) concentrations.

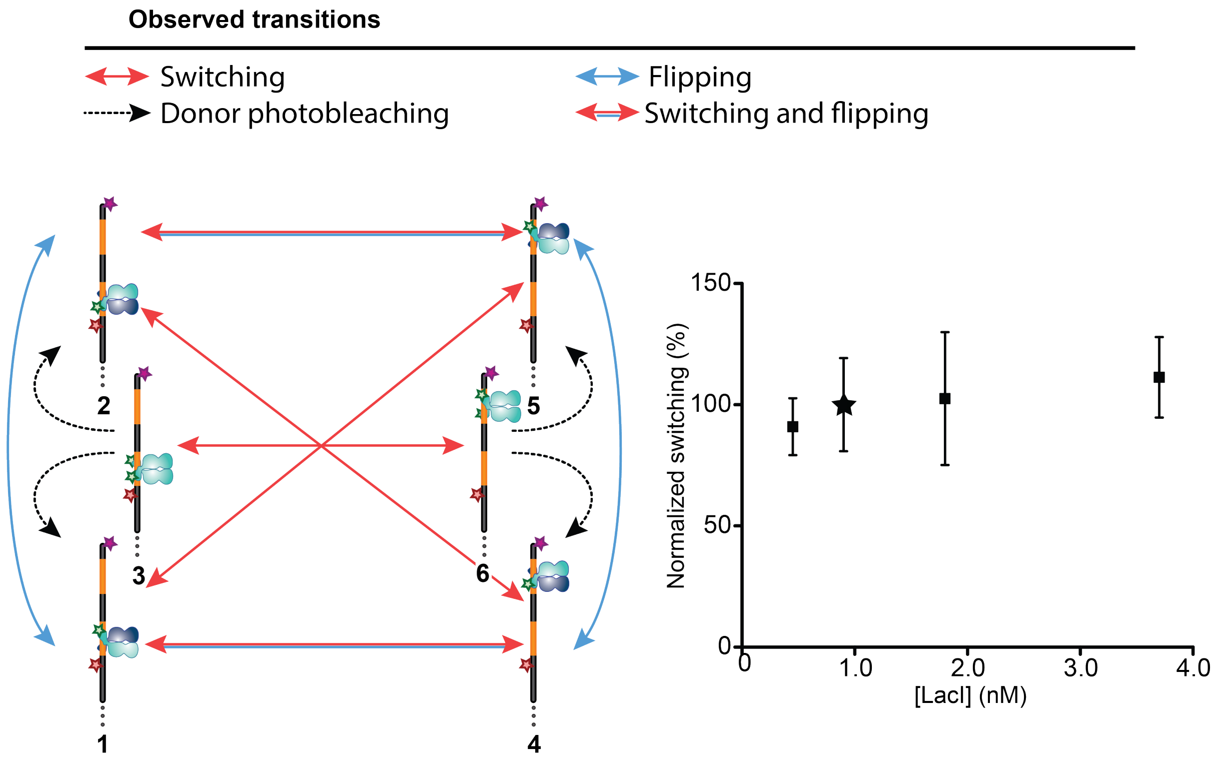


**Extended Data Figure 3. Predominant switching and flipping transitions.** Left: Cartoon schematic of the six distinct LacI-R binding states (see Methods) observed with a construct featuring two outer $O_1$ sites. Transitions between distinct states are depicted by arrows. Right: Switching rates observed at various LacI-R concentrations, normalized to the LacI-R concentration of 0.9 nM, (asterisk, used for the determination of all switching rates shown in Fig. 2) which is in the concentration regime where switching is not affected by the occupancy of multiple LacI molecules. Error estimates represent standard errors of the mean ($N$ > 100).

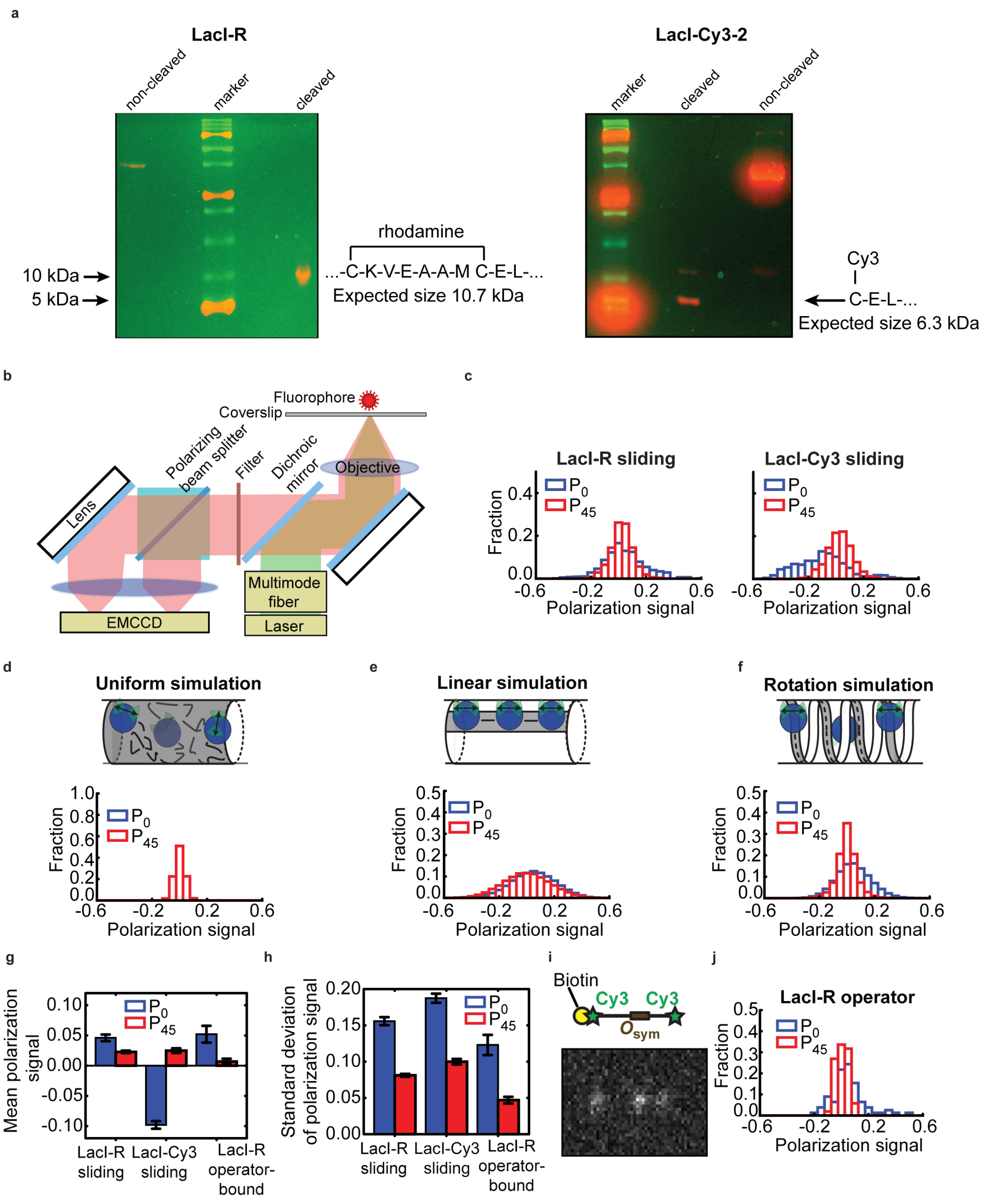


**Extended Data Figure 4. Camera-based polarization measurements and characterisation of dye labelling.** (**a**) SDS-PAGE analysis of LacI-R and LacI-Cy3-2 after peptide cleavage with CNBr. LacI-R was designed for bifunctional labelling (rhodamine attachment to both adjacent Cys residues) and LacI-Cy3-2 for monofunctional (using only one of the two Cys) labelling of the same α-helix. Precision Plus Protein™ Dual Xtra Standards (Bio-Rad) was used as ladder for both gels. (**b**) Schematic illustration of the experimental setup. (**c**) $P_0$ (blue) and $P_{45}$ (red) polarization distributions averaged over 600 ms (3 frames) per count measured for sliding LacI-R (left; 79 and 172 sliding events for $P_0$

and $P_{45}$, respectively) and LacI-Cy3 (right; 61 and 53 sliding events for $P_0$ and $P_{45}$, respectively). For $P_0$ measurements, the horizontal polarization axis was aligned with the stretching direction of DNA, whereas for $P_{45}$ measurements the horizontal polarization axis was rotated 45° away from the stretching direction of DNA (see Supplementary Information 4.2 for details). (**d**)-(**f**) Simulated polarization distributions for the uniform (**d**), linear (**e**), and rotation-coupled (**f**) sliding models. (**g**)-(**h**) Averages (**g**) and standard deviations (**h**) of experimental polarization distributions. Error bars represent standard errors. (**i**) Schematic illustration of the 7 kb DNA used in operator binding polarization measurements (top) and EMCCD image of LacI-R bound to the artificial, strong $O_{sym}$ operator on the flow-stretched DNA. (**j**) $P_0$ (blue) and $P_{45}$ (red) polarization distributions averaged over 600 ms (3 frames) per count measured for operator-bound LacI-R. Eleven and 15 binding events were detected for measurements of $P_0$ and $P_{45}$, respectively.

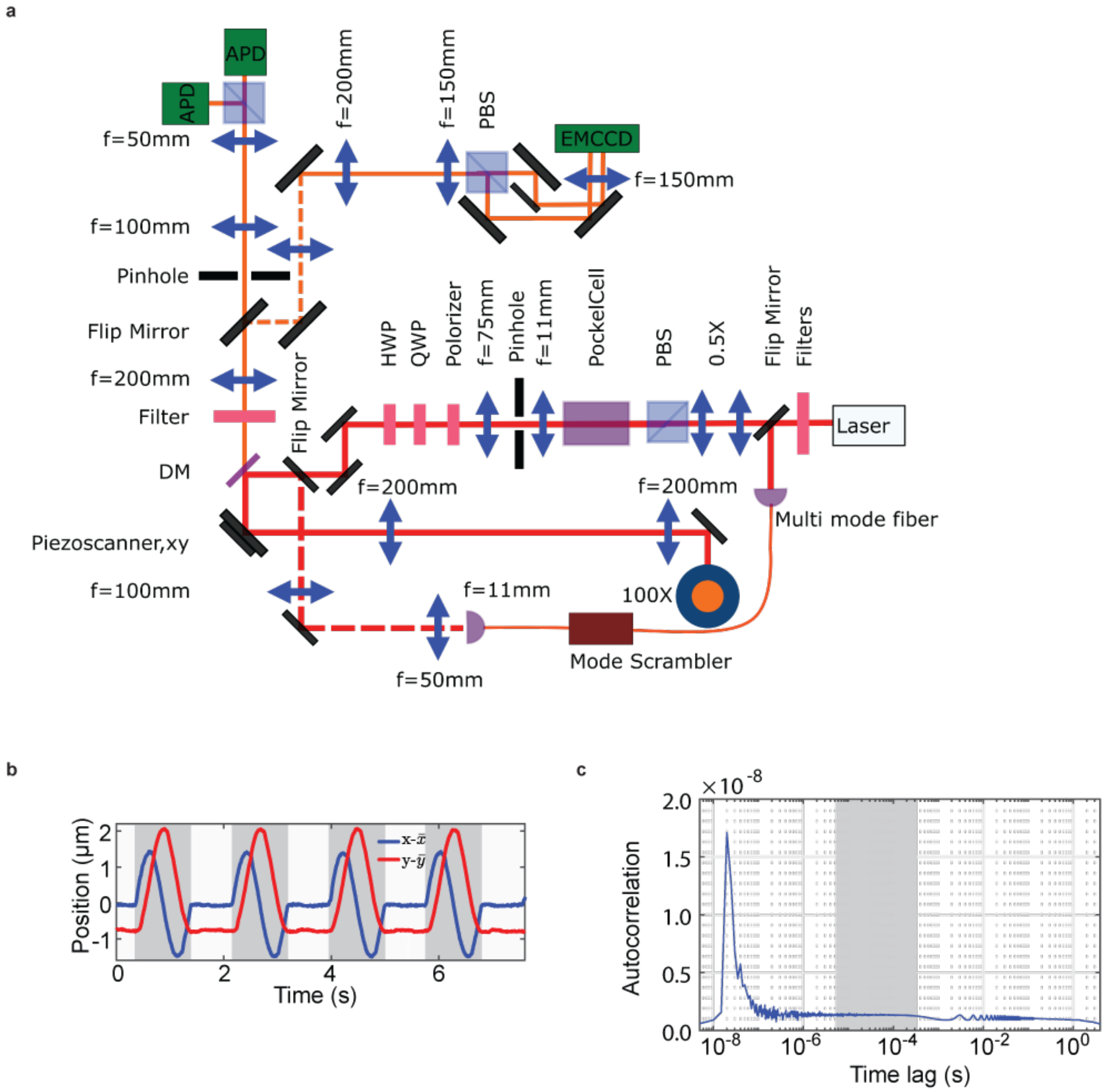


**Extended Data Figure 5. Optical layout and calibration data for SMCT-FCS.** (**a**) Optical layout of the combined system for fluorescence polarization microscopy and single-molecule confocal tracking. In the excitation path, a half-wave plate (HWP) and a quarter-wave plate (QWP) are placed behind a polarizer to create circularly polarized light at the objective. Abbreviations: polarization beam splitter (PBS), avalanche photodiode (APD), electron multiplying charge-coupled device (EMCCD), dichroic mirror (DM). (**b**) As a platform for testing the tracking capabilities of the real-time tracking system, immobilized fluorescent beads were used. Here, a tracking trajectory is shown where the stage is moving in circles with a diameter of 2.8 μm and the photon count rate is on average 24 kc/s (18 kc/s including laser off-time). Each greyed-out area represents a 1-s revolution followed by an 800 ms pause. In each paused section, the positioning standard deviation is 20 nm and includes stage noise from its feedback loop. (**c**) Autocorrelation function for a representative trajectory. Extraction of photon time tagging information and calculation of the autocorrelation. The plot depicts the high after pulsing peak at 20 ns, which rapidly decreases and flattens out after 500 ns. The grey area corresponds to the time regime plotted for autocorrelation functions in

Fig 3. After 1 ms an oscillation appears which is caused by the 4 ms instrument tracking period. The autocorrelation curve is here compensated for the 166 µs measurement off time between each 500 µs measurement.

**a**

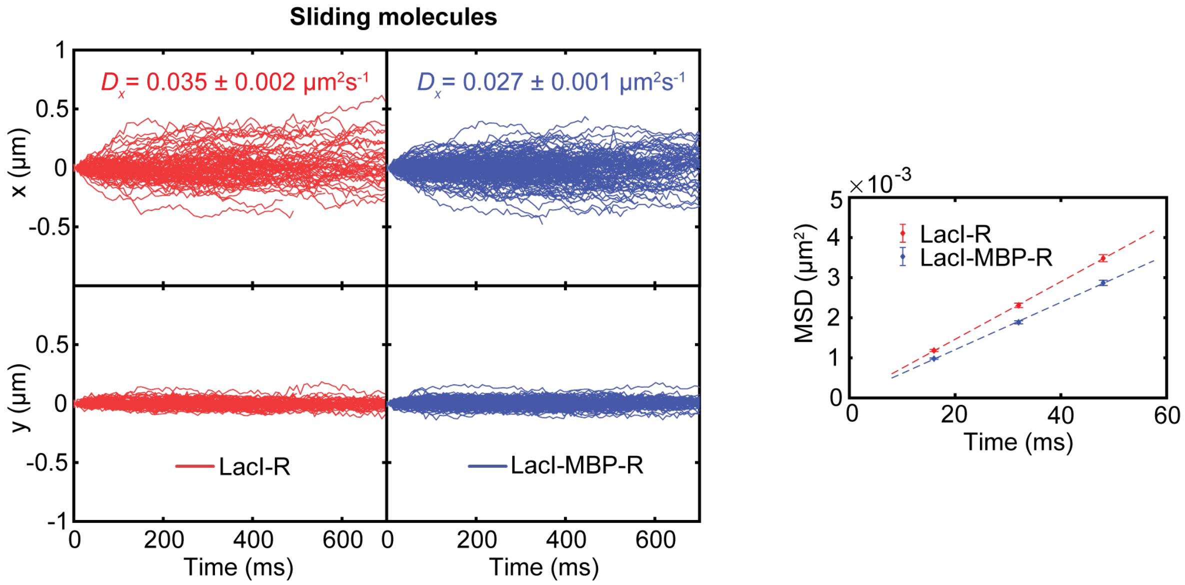


**b**

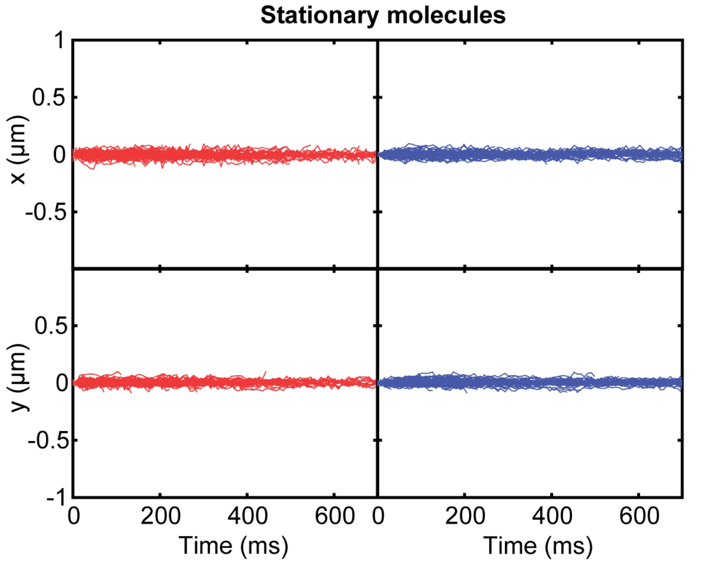


**Extended Data Figure 6. Traces captured with confocal tracking.** (**a**) (left) x (DNA direction) and y coordinate of sliding LacI-MBP-R (blue) and LacI-R (red) molecules obtained by confocal tracking. In total, 151 and 91 sliding molecules were captured for LacI-MBP-R and LacI-R, respectively. The average diffusion constants along the x-coordinate were 0.027 ± 0.001 $\mu m^2 s^{-1}$ and 0.035 ± 0.002 $\mu m^2 s^{-1}$. Diffusion constants are indicated as 20% trimmed means ± SEM. (right) Mean squared displacement for different time steps for LacI-MBP-R (bue) and LacI-R (red). (**b**) x (DNA direction) and y coordinate of stationary LacI-MBP-R (blue) and LacI-R (red) molecules obtained by confocal tracking. For clarity, 100 representative trajectories are shown for each LacI species.

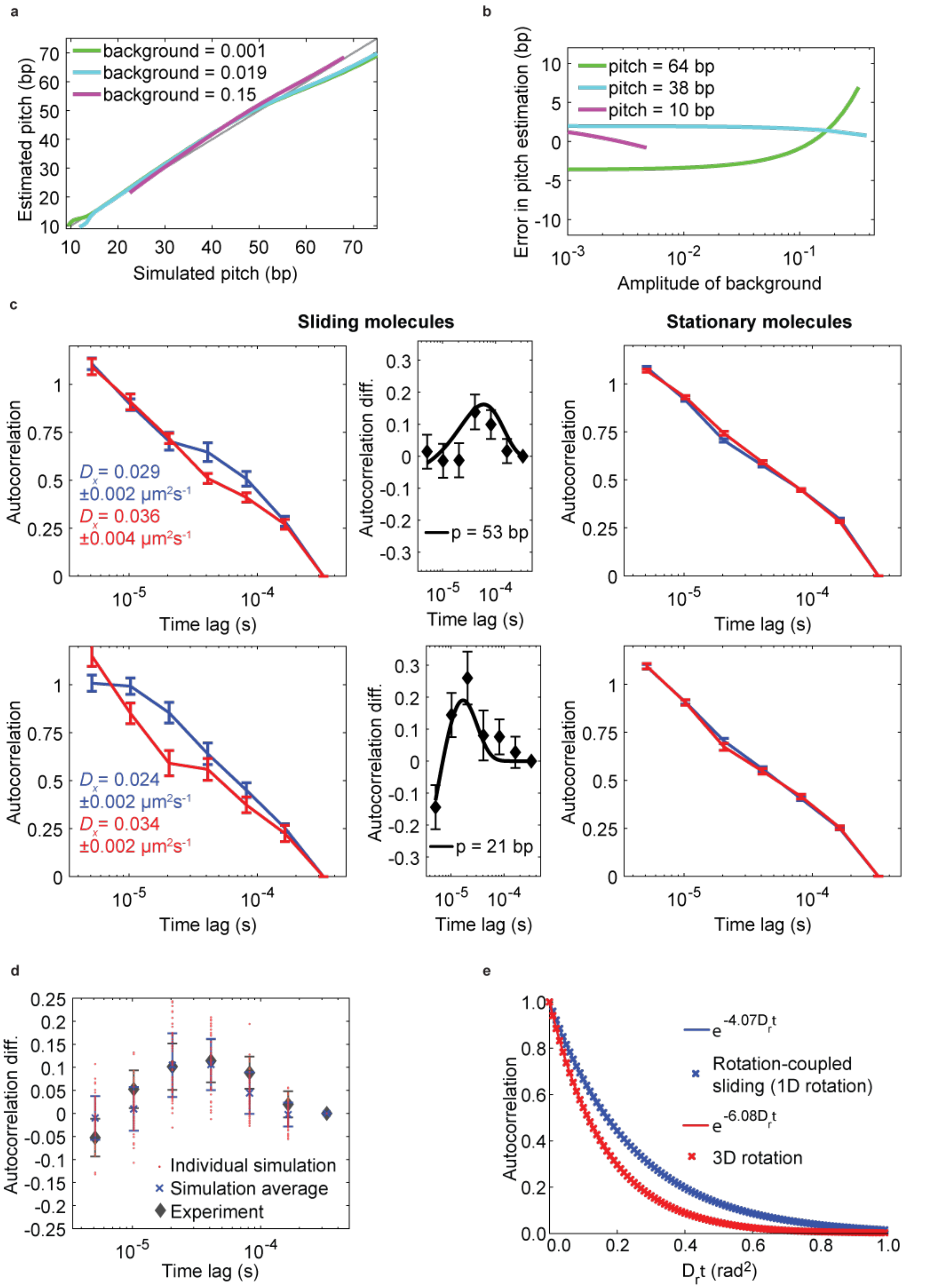


**Extended Data Figure 7. Fit of pitch-dependent autocorrelation model and repeats of SMCT-FCS experiments.** (**a**) Pitch estimates from simulated autocorrelations as a function of the pitch used in the simulation for different amplitudes of the background. (**b**) Error in the pitch estimate as a function of background amplitude for different pitches. To only consider simulated results relevant for the experimental results observed in Fig. 3, values are only reported for estimated pitches between 9 and 75 bp and when the largest distance (amplitude) of the difference in the simulated autocorrelation is at least 50% of the amplitude of the observed experimental difference. (**c**) The average normalized autocorrelation of the fluorescence signal for sliding (left) and stationary (right) LacI-MBP-R (blue) and LacI-R (red) molecules when the data has been pooled in two series (separated chronologically when the data was captured) for earlier (top) and later (bottom) recorded data. The reported diffusion constants are averages of all tracked molecules for each series. Error bars and error estimates represent standard errors. Diffusion constants are indicated as 20% trimmed means ± SEM .For the first experimental series 69 sliding and 2758 stationary molecules were analysed for LacI-MBP-R, and 36 sliding and 903 stationary molecules were analysed for LacI-R. For the second experimental series 82 sliding and 1015 stationary molecules were analysed for LacI-MBP-R, and 54 sliding and 691 stationary molecules were analysed for LacI-R. (middle) The difference in autocorrelation between LacI-MBP-R and LacI-R. Error bars represent standard errors. Lines represent the best fit to a rotation-coupled sliding model. (**d**) Difference in autocorrelation from simulations replicating the experimental data sets and filtering steps. The autocorrelation differences are indicated by the red dots. The number of traces, average trace length and average photon counts were the same as in the experiments, and the rotational diffusion constants were the fitted values from the experiments. The mean and expected standard error (blue crosses) was calculated as the mean and standard deviation of 60 individual simulations. The mean and standard error of the experiments (black diamonds) were estimated as described in Fig. 3g of the main text. (**e**) Autocorrelation functions calculated from intensities traces obtained with the vector-based dipole point spread function model without shot-noise. The dipole vector is rotated in Brownian dynamics simulations with unconstrained 3D diffusion (red crosses) or constrained diffusion around the x-axis (blue crosses). Lines represent best fits to single exponential functions of the autocorrelation data.

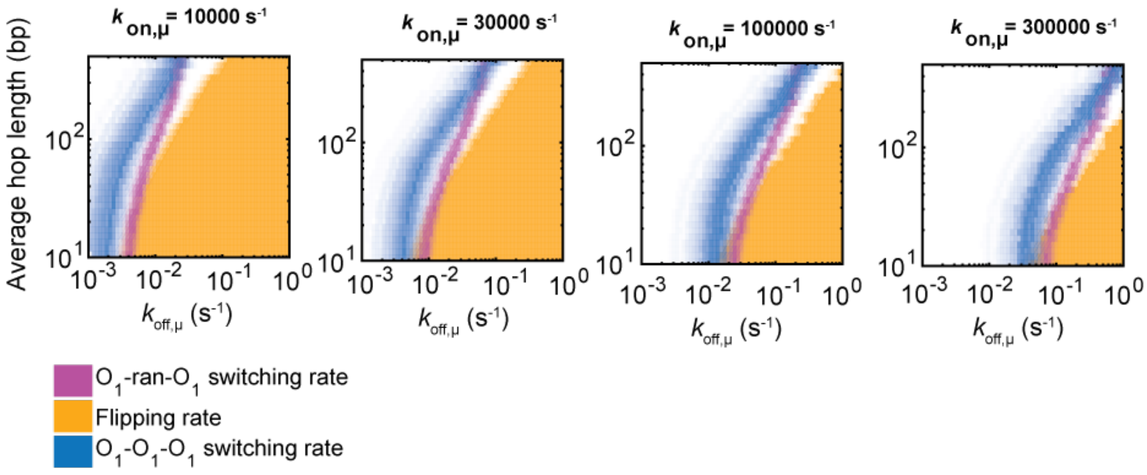


**Extended Data Figure 8. Probability densities of model parameters when taking into account the experimental FRET results.** Model parameters from simulations compatible with the observed rates for switching between two outer $O_1$ operators with (blue) and without (purple) an intervening internal $O_1$ operator, and compatible with the flipping orientation on the same operator (orange). The transparency values of the surfaces have been scaled with the probability density for the parameters given the experimental data. The mean and standard error of the upper bound of the average hop length calculated from the joint probability densities for the four different $k_{on,\mu}$ values are, from left to right: 22 ± 16 bp, 18 ± 11 bp, 16 ± 8 bp and 18 ± 12 bp.

**Extended Data Table 1. Sequences of LacI and single-molecule FRET DNA constructs.**

| Name | Sequence |
|---|---|
| LacI-R | MKPVTLYDVAEYAGVSYQTVSRVVNQASHVSAKTRCKVEAAMCELNYIPNR<br>VAQQLAGKQSLLIGVATSSLALHAPSQIVAAIKSRADQLGASVVVSMVERSG<br>VEAAKAAVHNLLAQRVSGLIINYPLDDQDAIAVEAAATNVPALFLDVSDQTPIN<br>SIIFSHEDGTRLGVEHLVALGHQQIALLAGPLSSVSARLRLAGWHKYLTRNQI<br>QPIAEREGDWSAMSGFQQTMQMLNEGIVPTAMLVANDQMALGAMRAITES<br>GLRVGADISVVGYDDTEDSSCYIPPLTTIKQDFRLLGQTSVDRLLQLSQGQAV<br>KGNQLLPVSLVKRKTTLAPNTQTHHHHHHG |
| LacI-Cy3 | MKPVTLYDVAEYAGVSYQTVSRVVNQACHVSAKTREKVEAAMAELNYIPNR<br>VAQQLAGKQSLLIGVATSSLALHAPSQIVAAIKSRADQLGASVVVSMVERSG<br>VEAAKAAVHNLLAQRVSGLIINYPLDDQDAIAVEAAATNVPALFLDVSDQTPIN<br>SIIFSHEDGTRLGVEHLVALGHQQIALLAGPLSSVSARLRLAGWHKYLTRNQI<br>QPIAEREGDWSAMSGFQQTMQMLNEGIVPTAMLVANDQMALGAMRAITES<br>GLRVGADISVVGYDDTEDSSCYIPPLTTIKQDFRLLGQTSVDRLLQLSQGQAV<br>KGNQLLPVSLVKRKTTLAPNTQTHHHHHH |
| LacI-MBP-R | MKPVTLYDVAEYAGVSYQTVSRVVNQASHVSAKTRCKVEAAMCELNYIPNR<br>VAQQLAGKQSLLIGVATSSLALHAPSQIVAAIKSRADQLGASVVVSMVERSG<br>VEAAKAAVHNLLAQRVSGLIINYPLDDQDAIAVEAAATNVPALFLDVSDQTPIN<br>SIIFSHEDGTRLGVEHLVALGHQQIALLAGPLSSVSARLRLAGWHKYLTRNQI<br>QPIAEREGDWSAMSGFQQTMQMLNEGIVPTAMLVANDQMALGAMRAITES<br>GLRVGADISVVGYDDTEDSSCYIPPLTTIKQDFRLLGQTSVDRLLQLSQGQAV<br>KGNQLLPVSLVKRKTTLAPNTQTGSGSGSEEGKLVIWINGDKGYNGLAEVG<br>KKFEKDTGIKVTVEHPDKLEEKFPQVAATGDGPDIIFWAHDRFGGYAQSGLL<br>AEITPDKAFQDKLYPFTWDAVRYNGKLIAYPIAVEALSLIYNKDLLPNPPKTWE<br>EIPALDKELKAKGKSALMFNLQEPYFTWPLIAADGGYAFKYENGKYDIKDVG<br>VDNAGAKAGLTFLVDLIKNKHMNADTDYSIAEAAFNKGETAMTINGPWAWSN<br>IDTSKVNYGVTVLPTFKGQPSKPFVGVLSAGINAASPNKELAKEFLENYLLTD<br>EGLEAVNKDKPLGAVALKSYEEELAKDPRIAATMENAQKGEIMPNIPQMSAF<br>WYAVRTAVINAASGRQTVDEALKDAQTNSSSHHHHHH |
| LacI-Far | MKPVTLYDVAEYAGVSYQTVSRVVNQASHVSAKTREKVEAAMAELNYIPNR<br>VAQQLAGKQSLLIGVATSSLALHAPSQIVAAIKSRADQLGASVVVSMVERSG<br>VEAAKAAVHNLLAQRVSGLIINYPLDDQDAIAVEAAATNVPALFLDVSDQTPIN<br>SIIFSHEDGTRLGVEHLVALGHQQIALLAGPLSSVSARLRLAGWHKYLTRNQI<br>QPIAEREGDWSAMSGFQQTMQMLNEGIVPTAMLVANDQMALGAMRAITES<br>GLRVGADISVVGYDDTEDSSCYIPPLTTIKQDFRLLGQTSVDRLLQLSQGQAV<br>KGNQLLPVSLVKRKTTLAPNTQTHHHHHHC |
| LacI-Cy3-2 | MKPVTLYDVAEYAGVSYQTVSRVVNQASHVSAKTREKVEAAMCELNYIPNR<br>VAQQLAGKQSLLIGVATSSLALHAPSQIVAAIKSRADQLGASVVVSMVERSG<br>VEAAKAAVHNLLAQRVSGLIINYPLDDQDAIAVEAAATNVPALFLDVSDQTPIN<br>SIIFSHEDGTRLGVEHLVALGHQQIALLAGPLSSVSARLRLAGWHKYLTRNQI<br>QPIAEREGDWSAMSGFQQTMQMLNEGIVPTAMLVANDQMALGAMRAITES<br>GLRVGADISVVGYDDTEDSSCYIPPLTTIKQDFRLLGQTSVDRLLQLSQGQAV<br>KGNQLLPVSLVKRKTTLAPNTQTHHHHHHG |
| Single $O_1$ site construct | Top strand:<br>5'-/5BioTinTEG/TCGTACTTCAAGTTTTGGGCGTGTCAAGTCCAAGGATTGC<br>TCTGTATACTTAAAAACGACGTGGCAGTAAAGGGAACGCAAGACTCTCAA<br>TCGCAATTGTTATCCGCTCACAATTCCGAAAGCCT-3'<br><br>Bottom strand:<br>5'-AGGCT/iCy5/TCGGAATTGTGAGCGGATAACAATTGCGAATGAGAGTCT<br>TGCGTTCCCTTTACTGCCACGTCGTTTTTAAGTATACAGAGCAATCCTTG |

| | |
|---|---|
| | GACTTGACACGCCCAAAACTTGAAGTACGA-3’ |
| Random sequence construct without $O_1$ site | Top strand:<br>5’-/5BioTinTEG/TCGTACTTCAAGTTTTGGGCGTGTCAAGTCCAAGGATTGCTCTGTATACTTAAAAACGACGTGGCAGTAAAGGAACGCAAGACTCTCA/iCy5/TCGCGATTGCAGCTCGAAGCAGCATCCGAAAGCC-3’<br><br>Bottom strand:<br>5’-GGCTTTCGGATGCTGCTTCGAGCTGCAATCGCGAATGAGAGTCTTGCGTTCCCTTTACTGCCACGTCGTTTTTAAGTATACAGAGCAATCCTTGGACTTGACACGCCCAAAACTTGAAGTACGA-3’ |
| $O_1$-*ran*-$O_1$ | Top strand:<br>5’-/5BioTinTEG/TCGTACTTCAAGTTTTGGGCGTGTCAAGTCCAAGGATTGCTCTGTATACTTAAAAACGACGTGGCAGTAAAGGAACGCAAGACTCTCA/iCy5/TCGCAATTGTTATCCGCTCACAATTCCGACAGCCAGGATTCCATGCCTCATGTGAAATTGTTATCCGCTCACAATTCGATAGCACATATCTAGCGATAT-3’<br><br>Bottom strand:<br>5’-ATATCGCTAGATATGTGC/iAlex750N/ATCGAATTGTGAGCGGATAACAATTTCACATGAGGCATGGAATCCTGGCTGTCGGAATTGTGAGCGGATAACAATTGCGAATGAGAGTCTTGCGTTCCCTTTACTGCCACGTCGTTTTTAAGTATACAGAGCAATCCTTGGACTTGACACGCCCAAAACTTGAAGTACGA-3’ |
| $O_1$-$O_1$-$O_1$ | Top strand:<br>5’-/5BioTinTEG/TCGTACTTCAAGTTTTGGGCGTGTCAAGTCCAAGGATTGCTCTGTATACTTAAAAACGACGTGGCAGTAAAGGAACGCAAGACTCTCA/iCy5/TCGCAATTGTTATCCGCTCACAATTCCGACAATTGTTATCCGCTCACAATTGTGAAATTGTTATCCGCTCACAATTCGATAGCACATATCTAGCGATAT-3’<br><br>Bottom strand:<br>5’-ATATCGCTAGATATGTGC/iAlex750N/ATCGAATTGTGAGCGGATAACAATTTCACAATTGTGAGCGGATAACAATTGTCGGAATTGTGAGCGGATAACAATTGCGAATGAGAGTCTTGCGTTCCCTTTACTGCCACGTCGTTTTTAAGTATACAGAGCAATCCTTGGACTTGACACGCCCAAAACTTGAAGTACGA-3’ |
| $O_1$-$O_3$-$O_1$ | Top strand:<br>5’-/5BioTinTEG/TCGTACTTCAAGTTTTGGGCGTGTCAAGTCCAAGGATTGCTCTGTATACTTAAAAACGACGTGGCAGTAAAGGAACGCAAGACTCTCA/iCy5/TCGCAATTGTTATCCGCTCACAATTCCGACGGCAGTGAGCGCAACGCAATTGTGAAATTGTTATCCGCTCACAATTCGATAGCACATATCTAGCGATAT-3’<br><br>Bottom strand:<br>5’-ATATCGCTAGATATGTGC/iAlex750N/ATCGAATTGTGAGCGGATAACAATTTCACAATTGCGTTGCGCTCACTGCCGTCGGAATTGTGAGCGGATAACAATTGCGAATGAGAGTCTTGCGTTCCCTTTACTGCCACGTCGTTTTTAAGTATACAGAGCAATCCTTGGACTTGACACGCCCAAAACTTGAAGTACGA-3’ |

Cysteine residues introduced for labelling are marked in red.
Modifications are reported using Integrated DNA Technologies (IDT) nomenclature. Naturally occurring LacI target sites $O_1$ and $O_3$ are highlighted in orange and brown, respectively.

**Extended Data Table 2. Sliding events in EMMCD and confocal tracking experiments.** Molecules are classified as sliding by our data analysis pipeline; sliding events in negative control measurements without $\lambda$-DNA thus reflect molecules that are falsely classified as sliding. A single-molecule detection event is defined as a captured trajectory that lasts for at least 600 (camera-based) or 200 (confocal tracking) ms. When no sliding events were detected, the upper limit for the last columns in the table have been calculated by assuming a single sliding event.

| **Experiment** | **Number of single-molecule detections** | **Number of sliding events** | **Sliding event per single-molecule detection** |
|---|---|---|---|
| LacI-R, 300 pM $\lambda$-DNA, camera tracking | 7782 | 251 | 0.032 |
| LacI-R, (-) $\lambda$-DNA, camera tracking | 968 | 2 | 0.0021 |
| LacI-MBP-R, 300 pM $\lambda$-DNA, confocal tracking | 8780 | 151 | 0.0172 |
| LacI-MBP-R, (-) $\lambda$-DNA, confocal tracking | 294 | 0 | <0.0034 |
