## Supplementary Information for "Mechanism of DNA surface exploration and operator bypassing during target search"

Emil Marklund, Brad van Oosten, Guanzhong Mao, Elias Amselem, Kalle Kipper, Anton Sabantsev, Andrew Emmerich, Daniel Globisch, Xuan Zheng, Laura Lehmann, Otto Berg, Magnus Johansson, Johan Elf* & Sebastian Deindl*

#### 1 Protein expression, purification and labeling

For LacI-R and LacI-MBP-R, C-terminally 6xHis-tagged variants of LacI were expressed from pD861 plasmids in BL21 *E. coli* cells. 20 ml overnight cultures were diluted into 2 L LB medium with 50 µg/ml kanamycin and grown at 37 °C and 120 RPM agitation. The expression of LacI was induced at $OD_{600} = 0.5$-1 by adding L-rhamnose to a final concentration of 0.2 % (volume/weight). After 3 hours, the cells were harvested by centrifugation at 6,000 x g at room temperature for 40 min. The pelleted cells were then resuspended in 20 mL buffer A (500 mM NaCl, 20 mM imidazole, 5 mM 2-mercaptoethanol, 20 mM phosphate, pH 7.4) containing 1 EDTA-free protease inhibitor pill (Roche) and lysed on ice for 40 min after the addition of lysozyme and benzonase I to final concentrations of 10 µg/ml and 25 units/mL, respectively. After additional lysis using a french press, the lysate was clarified by centrifugation at 3800 xg for 1 hour followed by filtering of the supernatant through a 0.45 µm cellulose acetate membrane filter. Purification was carried out at 4 °C and a flow rate of 1 mL/min on an ÄKTA pure 25 FPLC system (GE Healthcare) where the lysate was applied to a 5 mL HisTrap HP affinity purification column (GE Healthcare) equilibrated with buffer A. After washing with 10 column volumes of buffer A, bound protein was eluted with a linear imidazole gradient (20-500 mM) and collected in 3 ml fractions. Fractions were analysed with SDS-PAGE and the LacI-containing fractions were pooled and buffer exchanged using 10 kDa cut-off Amicon Ultra-15 Centrifugal filters (Merck Millipore) into phosphate-buffered saline (PBS) supplemented with 10 % glycerol.

Labeling of LacI with N,N´-Bis[2-(iodoacetamido)ethyl]-N,N´-dimethylrhodamine (Santa Cruz Biotechnology) was performed in PBS + 10 % glycerol. 12 mL of labeling buffer were first degassed at room temperature using vacuum for 30 min, followed by the addition of TCEP (10 mM in PBS + 10 % glycerol, pH adjusted to 7.4), and purified LacI to final concentrations of 400 µM and 40 µM, respectively. After incubation at room temperature for 30 min, the dye (1 mM dissolved in PBS) was added to the labeling reaction to a final concentration of 60 µM. The labeling reaction was incubated in the dark and at room temperature with light agitation for 90 min, followed by quenching with the addition of 2-mercaptoethanol to a final concentration of 5 mM. The quenched reaction was prepared for removal of unreacted dye by a 1:1 dilution in buffer A, followed by application to a 5 mL HisTrap HP affinity purification column equilibrated with buffer A at 4 °C. After washing with 10 column volumes of buffer A for removal of unreacted dye, bound protein was eluted with buffer A supplemented with an imidazole concentration of 500 mM and collected in 1.5 ml fractions. Fractions were analyzed with SDS-PAGE and visualized on a ChemiDoc MP (BioRad) using green epi LED excitation light (Extended Data Fig. 1b). The eluted labeled fractions were pooled and buffer exchanged into PBS supplemented with NaCl 500 mM using a 10 kDa cut-off Amicon Ultra-15 centrifugal filter. Finally, labeled protein with a dye concentration of $\sim$ 50 µM was diluted 1:1 with 100% glycerol, flash frozen in liquid nitrogen, and stored at -80 °C for later use.

After labeling with bifunctional rhodamine, the majority of the labeled protein ran as a monomeric band when separated on a SDS-PAGE gel, while a minor fraction of the protein ran as a band corresponding to a dimer (Extended Data Fig. 1b). The fact that a minor fraction of LacI subunits persisted as dimers even after denaturation could be the result of inter-subunit crosslinking by the bifunctional rhodamine dye. We note however that such covalently crosslinked LacI dimers are unlikely to affect our single-molecule experiments for two reasons. First, they are stoichiometrically outnumbered by a vast excess of non-crosslinked, dye-labeled LacI forming the naturally occurring dimer. Second, the specific location on the DNA-binding domain of the cysteine residues used for labeling LacI implies that a potential dye-mediated crosslink between two monomers would give rise to a completely unphysiological situation with a structure that dramatically deviates from the DNA-binding competent LacI dimer. Thus, structural considerations make it extremely unlikely that the small fraction of covalently crosslinked LacI could bind to DNA and contribute to the single-molecule experiments. Furthermore, the dimeric band was also present when LacI was labeled at a single cystein further away from the DNA binding domain with a Cy3 fluorophore with only one reactive group (LacI-Far, Extended Data Fig. 1b). This indicates that the dimeric band on the gels are caused by non-crosslinked correctly labeled dimers that fail to denature, and not only by LacI dimers that are crosslinked by the fluorophore. Additionaly, our peptide cleavage assay with CNBr yield a distinct band around 10 kDa for LacI-R (consistent with correct bifunctional labelling), but no bands corresponding to higher molecular weights as would be expected after cleaving inter-LacI crosslinked species (Extended Data Fig 4j), further indicating that no covalently crosslinked LacI is present in the sample.

LacI-Far, LacI-Cy3 and LacI-Cy3-2 was expressed, purified and labeled according to previously published methods[1]

### 2 Flow channel preparation for single-molecule tracking

#### 2.1 Single-molecule sliding assay

Preparation of flow channels with tethered $\lambda$-DNA (New England Biolabs) was carried out according to previously published methods[2]. Biotinylated coverslips were assembled into flow channels with 3 mm x 0.0089 mm cross-sectional areas. After assembly, flow channels were incubated with 1 mg/ml Neutravidin (Thermo Scientific), 1 mg/ml bovine serum albumine (BSA) dissolved in 50 mM Tris buffered at pH 7.5 supplemented with 10 mM NaCl for 5 minutes. The following steps, including single-molecule microscopy, were performed in an imaging buffer containing 10 mM potassium phosphate pH 7, 1 mM NaCl, 0.1 mM EDTA, 5% (v/v) glycerol, 0.5 mg/ml BSA and 1 mM β-mercaptoethanol. To reduce fluorophore photo-bleaching and blinking, the imaging buffer was degassed and supplemented with 1 mM Trolox prior to use. The Neutravidin solution was replaced by buffer containing 300 pM $\lambda$-DNA biotinylated on one end for 1 hour in order to tether one end of the DNA to the cover slip. All single-molecule sliding experiments were then performed with DNA stretched with a flow rate of 1 ml /min with fluorescent protein concentrations of 10-100 pM.

#### 2.2 Single-molecule operator binding assay

A 7 kb dsDNA with a Cy3 fluorophore on each end, biotin on one end, and an internal $O_{sym}$ operator site 4 kb into the sequence was generated by PCR in two steps. In both PCR steps, amplification was done with Phusion Hot Start II DNA polymerase (Thermo Scientific) in PCR mixtures supplemented with 3% DMSO. In the first step, a genomic DNA prep. of *E. coli* strain JE101[3] was used as template at a final concentration of 1 ng/µl to create an intermediate product, slightly longer than the final 7 kb product. In this step, an annealing temperature of 67°C with primers osym_F_v3 (5'-ACCAGCGAAGCATTGTGTT-3') and osym_R_v3 (5'-CGCCCTGGTTAGTTCAACAT-3') was used. The intermediate product was purified with the PureLink quick PCR purification kit (Invitrogen) before being used as the template for the second PCR reaction at a final concentration of 0.2 ng/µl. In the second PCR step, annealing temperatures between 50 and 56°C with primers osym_F_BIOCY3_v2 (5'-/5BiosG/TCGAGG/iCy3/AGTAGCGTTTTAC-3') and osym_R_CY3_v2 (5'-CACTAGC GTACGAATAGCG/iCy3/AACCTGGTATCAA-3') were used. To remove any additional undesirable PCR products, the PCR mixture was purified on a 0.8% agarose gel after the reaction. Bands of the expected size of 7 kb were excised from the gel and purified with the PureLink Quick Gel Extraction Kit (Invitrogen). Purified operator DNA was eluted in 10 mM Tris pH 7.5 and stored at -20°C for later use. All primers were ordered from IDT and designed so that internal Cy3 fluorophores replaced one nucleotide in the sequence complementary to the template.

The single-molecule operator binding assay was prepared in the same way (preparation of flow-channels and buffers etc.) as the single-molecule sliding assay unless otherwise stated. Operator DNA (1 nM) and fluorescent LacI (200 nM) were pre-incubated in imaging buffer for 1.5 h at room temperature prior to tethering of the operator DNA to the coverslip. The mixture of operator DNA/LacI was then diluted in 10 ml imaging buffer (0.15 pM/30 pM final concentration) and introduced into the flow channel with a flow rate of 1 ml/min. Single-molecule operator binding events were then observed in the imaging buffer with DNA stretched with a flow rate of 1 ml/min. When three fluorescent dots (Cy3-LacI-Cy3) with the correct spacing were found in the flow channel, movies were recorded for polarization analysis. The fluorescent dots were then validated to be on flow stretched DNA by turning off the flow (and observing the relaxation of the DNA) followed by turning on the flow again (and observing the DNA being stretched out, Extended Data Fig. 4i).

### 3 Microscopy

#### 3.1 Single-molecule Fluorescence Resonance Energy Transfer (sm-FRET)

The presence of two monomers within each LacI dimer gave rise to a heterogeneous population of donor-labelled LacI dimer with three different labelling configurations: (1) donor on the monomer subunit proximal to the acceptor, (2) donor on the monomer subunit distal to the acceptor, (3) donor on both LacI monomers. Single-molecule detection allowed these configurations to be discriminated. LacI containing a single donor dye on the distal or proximal monomer subunit gives rise to a histogram of mean FRET values for binding events with two distinct peaks centred at FRET = 0.16 and FRET = 0.89, respectively. The assignment of these peaks to these two labeling configurations was further confirmed by individual FRET time traces, which showed one-step photobleaching due to LacI dimer bearing a single donor dye. LacI with donor on both LacI monomers gave an intermediate FRET value of 0.60 and was often observed with two-step photobleaching, where the intermediate FRET transitioned into either the distal or proximal population before dissociating/bleaching.

##### 3.1.1 Analysis of single-molecule FRET time traces from single-site constructs

All time traces were manually groomed to include only binding events that contained regions with first a sudden appearance (LacI binding) of fluorescence signals and FRET followed by their sudden disappearance (LacI dissociation). Binding events which contained bleaching of the Cy5 dye were also removed from analysis. These Cy5 bleaching events were characterised by the disappearance of the Cy5 signal while the donor signal remained essentially unchanged. To groom the time traces, the edges of each region of interest were marked and all timepoints outside of these regions were considered to reflect the unbound state (set to a FRET value of -1 for clear distinction from other states in Hidden Markov Model (HMM) analysis), while the regions of interest were left unaltered. Full time traces devoid of any binding events were excluded from analysis. The groomed traces were concatenated and analyzed using the HMM.

##### 3.1.2 Analysis of single-molecule FRET time traces from multi-site constructs

The grooming of time traces for the three colour data from multi-site constructs was handled similar to the analysis of traces from single-site constructs to only include binding events with sudden appearance/disappearance of both donor and acceptor signals. The addition of multiple sites creates an ambiguous 'dark', zero-FRET state, where we observe a LacI molecule on the DNA but are unsure of its location. The lack of acceptor intensity in these dark regions can have several ambiguous explanations. Firstly, it could be that the acceptor dye has bleached. Secondly, it could be that a LacI molecule has transferred to the central intervening site without any acceptor fluorophore within FRET range. Lastly, it could be that the LacI molecule has switched to the second flanking site which has a bleached/missing fluorophore. Supplementary Figure 1a provides an example time trace containing a dark state which can be explained as a LacI molecule moving first from the top to the central intervening site and then to the bottom flanking site. Alternatively, the time trace could equally be explained by the fluorophore on the top site bleaching at t=393 s before LacI switches directly to the second, bottom site some time later, thereby overshooting the intervening site. Similarly, Supplementary Figure 1b shows time traces with a dark regime that may be interpreted as stemming from a LacI molecule shifting from the bottom site to the intervening site in the center and then back to the bottom site. We caution however that these traces are ambiguous in that they might alternatively be explained by the direct switching to a flanking site with an already bleached fluorophore. Due to this ambiguity, LacI binding to the intervening site cannot be assigned with certainty. We therefore removed all such traces (with a zero-FRET state) from analysis and indirectly inferred trapping by the intervening site from the change in switching rates between the unambiguously assigned flanking sites in our HMM analysis.

Alexa750 exhibits lower photostability than Cy5. Moreover, on our setup, we cannot directly excite Alexa750 to confirm its presence (as opposed to Cy5). Therefore, an unknown subset of multi-site DNA molecules lacks their Alexa750 dye, which leads to underestimating the frequency of bottom site (acceptor: Cy5) → top site (acceptor: Alexa750) switching transitions. To avoid this bias, we focused our HMM analyses on top (acceptor: Alexa750) → bottom site (acceptor: Cy5) switching transitions.

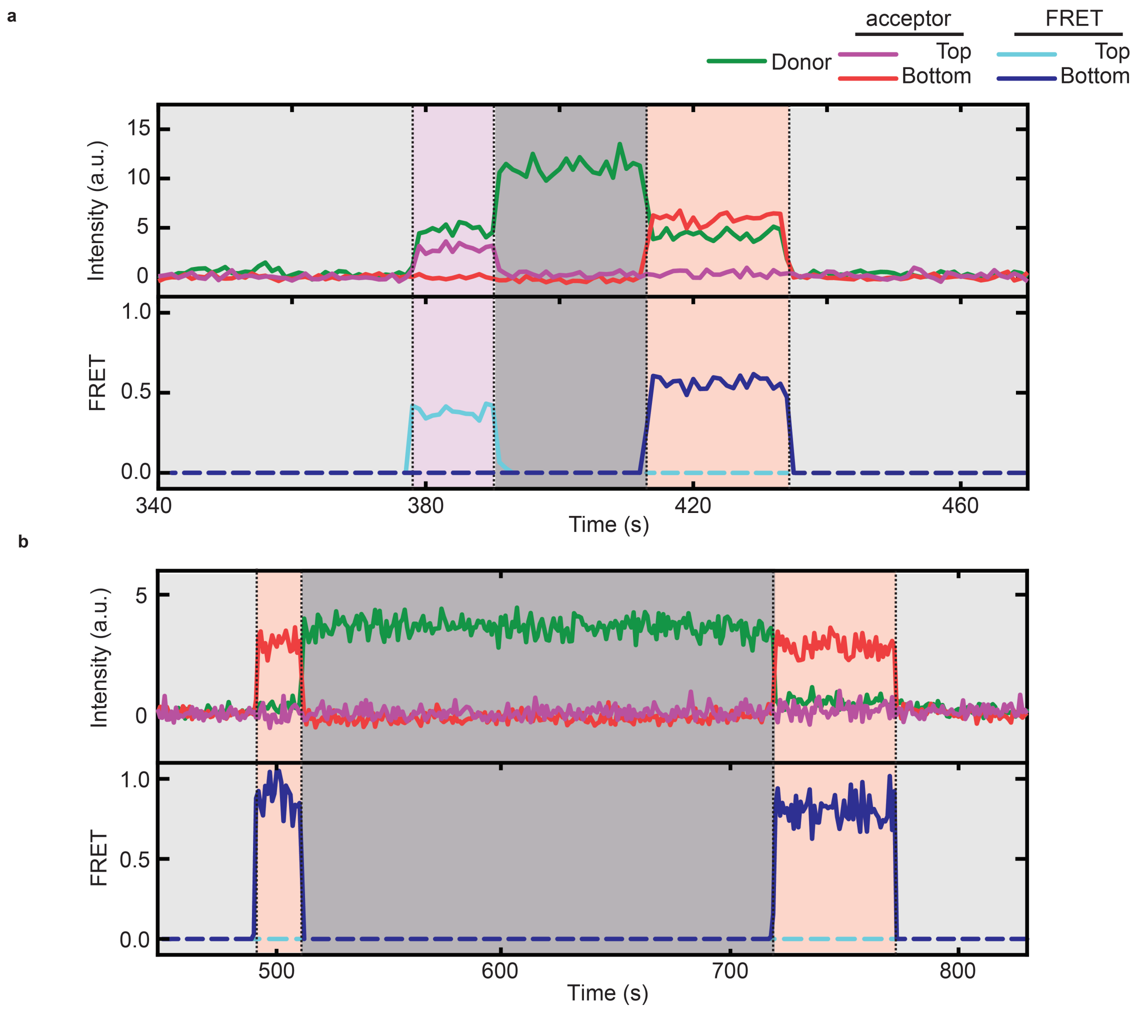


**Supplementary Figure 1. Sample time traces.** Donor fluorescence (green), bottom (red) and top (purple) site acceptor fluorescence, as well as FRET from the bottom (dark blue) and top site (light blue). (a) Example trace illustrating a 'dark' LacI state. Within the same binding event, a non-zero donor signal from the LacI molecule persists, while the acceptor signal first changes from a distinct value corresponding to the top Alexa750 site at t=378 s to the absence of any discernible signal in either acceptor channel at t=393s, to finally reach a distinct value in the Cy5 acceptor channel corresponding to LacI bound at the opposing bottom Cy5 site at t=412 s. (b) A second example of a 'dark' LacI state. In a single binding event, a non-zero donor signal from the LacI molecule persists, while the acceptor signal changes from a distinct value in the Cy5 channel (LacI bound at the bottom site site) at t=492 s to no discernible signal in either acceptor channel at t=514 s, followed by the reappearance of a Cy5 acceptor signal corresponding to LacI bound at the bottom site at t=718 s.

#### 3.2 Single-Molecule Confocal laser Tracking with Fluorescence Correlation Spectroscopy (SMCT-FCS)

##### 3.2.1 Real time tracking optimization

The confocal tracking with real time feedback was optimized for tracking with good stability for slow moving objects. For optimization we used fluorescent 100 nm beads which were immobilized on a microscope coverslip. The bead sample was placed on a xyz-piezo stage (Thorlabs Nano max) and imaged by the confocal system. The xyz-stage was programmed to move in circles with a diameter of 2.8 µm on the xy-plane. The revolution time was set to 1 second followed by a 800 ms pause before starting the next revolution (Extended Data Fig. 5b). Photon count rates for the bead tracking were on average 24 k counts/s (18 k counts/s including laser off time) which is about twice as high as the count rate of the fluorescently labeled LacI.

Since there are some intensity variations between experimental runs we performed a calibration before each sliding experiment. When calibrating, the sliding experiment was started together with a circular motion on the xy-stage. The tracking system parameters for entering and releasing the tracking mode were optimized using this easy to spot motion when the tracking system locked onto stationary fluorophores bound to the glass surface. Optimization of lock-in and release parameters was done by observing that the tracking was catching the rotational movement and releasing properly when the photon counts dropped below the mean photon threshold to avoid background driven tracking.

#### 3.3 Wide-field fluorescence polarization microscopy

The two polarization components emitted from the same fluorophore were first correlated to each other by translational registration of a 10 µm x 10 µm grid, imaged prior to each experiment. Single molecules were detected and localized using the sum of both polarization components. An à trous wavelet three-plane decomposition was used for dot detection[4].

Dots were detected through scale-dependent standard deviation thresholding in the third wavelet plane, where the standard deviation was estimated by the median absolute deviation method[5] and the threshold was set to 3 standard deviations. Dot centers were localized by calculating the weighted centroid from the pixel regions obtained from dot detection. Centroids in different frames of the movie were then connected into trajectories with u-track[6] using a maximal search radius of 4 pixels for the sliding experiments and 3 pixels for the operator binding experiments. To account for fluorophore blinking, a maximum dark time of 2 frames was used in the sliding experiment, while no upper limit was set on the dark time in the operator binding experiments (trajectory building in the operator experiments is more trivial since the dots do not move).

Classification of sliding trajectories was done with principal component analysis (PCA) on trajectories longer than 15 frames (3 s). Trajectories were only classified as sliding if their principle direction of movement did not deviate more than 20° from the expected DNA stretching direction (see Supplemental Table 1 for more classification parameters). The classifier was optimized to effectively find sliders in flow-channels with DNA but not in negative control channels without DNA (Extended Data Table 2).

In operator binding control experiments with stretched Cy3-labeled DNA, binding events were analyzed by selecting the trajectories corresponding to the LacI dot from the EMCCD data (located approximately 4 kb $\sim$ 13 pixels and 3 kb $\sim$ 10 pixels away in the DNA stretching direction from the 5'- and 3'- Cy3 dot, respectively).

The background intensities of pixels were estimated by calculating a moving average of each fluorescence image, implemented with exclusion of outliers to avoid including fluorescent dots in the average. The two polarization intensities of single molecules in each frame were calculated as the difference between the raw fluorescence intensity and the background, summed over a 7 x 7 pixel square around the centroid of the dot. After summation of the intensities over 3 frames (600 ms), the polarization signal was then calculated according to

$$P_\beta = \frac{I_\beta - I_{\beta+90}}{I_\beta + I_{\beta+90}} \tag{1}$$

The polarization signal for individual LacI molecules sliding on DNA was measured with one polarization axis aligned with the stretching direction of DNA (measuring $P_0$), as well as shifted 45° from it (measuring $P_{45}$). These camera-based polarization measurements can be used to rule out some modes of sliding. The rationale for the analysis is that the polarization signal is averaged out and thus close to zero in the directions corresponding to axes of symmetry where the protein is free to rotate. On the other hand a preferential orientation would manifest itself in a broadening of the distribution of polarizations. For example, if the protein was free to move and rotate on the DNA surface in all directions at a time scale faster than the imaging frame rate, the polarization distribution would be narrow in both $P_0$ and $P_{45}$. As a reference, this case was simulated using a vector-based point-spread function model for translating dipole orientation[7] (Extended Data Fig. 4d). In contrast, if the protein did not rotate around any of its own axes and only diffused linearly along the DNA any bias in direction of the DNA, protein or dye on the imaging time-scale would give rise to non-zero polarization signals resulting in a broadening in both $P_0$ and $P_{45}$ (Extended Data Fig. 4e). Furthermore, $P_0$ would have a non-zero average if the dye exhibited a preferred orientation with respect to DNA whereas $P_{45}$ would be indifferent with respect to DNA-protein alignment. The addition of a fast rotation of the protein around the DNA axis would not change $P_0$ compared to the linear diffusional sliding, whereas $P_{45}$ would be averaged out due to the symmetry of the rotation (Extended Data Fig. 4f). The experimental distributions for LacI labeled with rhodamine or Cy3 (LacI-R and LacI-Cy3) both display broadening in $P_0$ as compared with $P_{45}$. When compared with the simulations, this experimental result is only compatible with fluorophore rotation around the DNA axis.

To investigate the sources of noise in the measurement, we compared the polarization of the sliding LacI-R to that of operator bound LacI-R (Extended Data Fig. 4i,j). We found that their average $P_0$ polarization signals were very similar, suggesting that the protein may slide in an orientation that is close to the operator bound structure. The $P_0$ standard deviation was slightly smaller for the operator bound conformation, showing that only some of the broadening could be attributed to protein DNA interactions while sliding. Most importantly however, $P_{45}$ was averaged out also for the operator bound molecule suggesting that DNA itself was rotating around its own long axis on the imaging time scale.

#### 3.4 Sliding classification parameters

Table 1: **Classification and cut-off parameters in EMCCD experiments.**

| Parameter or property | Value |
|---|---|
| Sliding direction | Closer than 20° from expected DNA stretching direction |
| Minimum trajectory length | 3 s |
| Minimum eigenvalue of largest principal component (of molecule position with 200 ms frame rate) | 0.05 µm$^2$ |
| Maximum eigenvalue of smallest principal component (of molecule position with 200 ms frame rate) | 0.012 µm$^2$ |

Table 2: **Classification and cut-off parameters in confocal tracking experiments.**

| Parameter or property | Value |
|---|---|
| Sliding direction/ Direction of largest principal component (of molecule position) | Closer than 20° from expected DNA stretching direction |
| Minimum trajectory length | 0.55 s |
| Minimum eigenvalue of largest principal component (of molecule position) | 0.0015 µm$^2$ |
| Maximum eigenvalue of smallest principal component (of molecule position) | 0.00035 µm$^2$ |
| Maximum eigenvalue of largest principal component (of molecule position) in a 10 frame (80 ms) moving window | 0.016 µm$^2$ |
| Minimum eigenvalue of largest principal component (of molecule position) in a 10 frame (80 ms) moving window | 0.00008 µm$^2$ |
| Maximum diffusion coeff. along sliding direction | 0.07 µm$^2$ s$^{-1}$ |
| Minimum diffusion coeff. along sliding direction | -0.005 µm$^2$ s$^{-1}$ |

### 4 Models

#### 4.1 Vector based dipole point spread function model

The model used for the polarization dependent emission of a dipole are based on the work in[7]. Here the emitted electric field of a dipole $\bar{\mu}$ is described

$$\bar{\mu} = \begin{pmatrix} \sin(\Theta) \cdot \cos(\Phi) \\ \sin(\Theta) \cdot \sin(\Phi) \\ \cos(\Theta) \end{pmatrix} \quad (2)$$

which is parameterized by the two angels $\Theta \in [0, \pi]$ and $\Phi \in [0, 2\pi)$. At the back focal plane of the objective the electric field can be defined as $\mathbf{E}(\phi, \rho) = A \cdot \mathbf{G}_{bfp}(\phi, \rho) \cdot \bar{\mu}$ where

$$\mathbf{G}_{bfp}(\phi, \rho) = e^{i \cdot n \cdot k \cdot d \cdot \sqrt{1-\rho^2}} \cdot \frac{e^{i \cdot n \cdot k \cdot f_{obj}}}{4\pi \cdot f_{obj}} \sqrt{\frac{n}{n_{bfp}}} \cdot \frac{1}{(1-\rho^2)^{1/4}} \cdot \begin{pmatrix} \sin^2(\phi) + \cos^2(\phi) \cdot \sqrt{1-\rho^2} & \sin(2\phi) \cdot (\sqrt{1-\rho^2} - 1)/2 & -\rho \cdot \cos(\phi) \\ \sin(2\phi) \cdot (\sqrt{1-\rho^2} - 1)/2 & \cos^2(\phi) + \sin^2(\phi) \cdot \sqrt{1-\rho^2} & -\rho \cdot \sin(\phi) \\ 0 & 0 & 0 \end{pmatrix} \quad (3)$$

is the 3x3 Green tensor which converts the dipole far field to the back focal plane of the objective, $n$ and $n_{bfp}$ are the refractive index surrounding the dipole and at the back focal plane respectively, $k = 2\pi/\lambda$ is the wavenumber, $f_{obj}$ is the objective focal length which is related to the magnification $M = (n/n_{bfp}) \cdot (f_{tub}/f_{obj})$ and the tube lens $f_{tub}$, $d$ is the distance to the dipole and the pair $(\phi, \rho)$ is the parameterization of the back focal plane with the radius $\rho$ and angle $\phi$. The parameter $\rho$ is directly related to the numerical aperature through $NA = n \cdot \rho$. Here we assume that the dipole is situated on the glass interface of a microscope coverslip and not floating freely in a water solution, this allows us to set $n = 1.518$ which means $\rho \in [0, 0.96]$. The last row in the Green tensor $\mathbf{G}_{bfp}(\phi, \rho)$ are all zeros because there is no z component in the polarization field at the back focal plane, which is a consequence of the paraxial approximation used when deriving the Green tensor.

The field components in the back focal plane are then

$$\begin{pmatrix} X(\phi, \rho, \Theta, \Phi) \\ Y(\phi, \rho, \Theta, \Phi) \\ Z(\phi, \rho, \Theta, \Phi) \end{pmatrix} = A \cdot \mathbf{G}_{bfp}(\phi, \rho) \cdot \bar{\mu} \quad (4)$$

where $A$ is the amplitude of the dipole moment (which is dependent on the polarization of the excitation light and the dipole orientation, $A = 1$ for isotropic excitation light, $A = sin\Theta$ for excitation light circularly polarized in the xy-plane). If a polarization detector is placed at the image plane it will measure the total intensity over the whole point spread function. Since we are only interested in the polarization signal over the dipole we do not need to implement the tube lens but instead directly integrate each polarization intensity component over the back focal plane radially out to $\rho_{max} = NA/n$.

$$\begin{pmatrix} I_X(\rho_{max}, \Theta, \Phi) \\ I_Y(\rho_{max}, \Theta, \Phi) \\ I_z(\rho_{max}, \Theta, \Phi) \end{pmatrix} = \begin{pmatrix} \int_0^{2\pi} \int_0^{\rho_{max}} \left|X(\phi, \rho, \Theta, \Phi)\right|^2 \rho \cdot d\rho d\phi \\ \int_0^{2\pi} \int_0^{\rho_{max}} \left|Y(\phi, \rho, \Theta, \Phi)\right|^2 \rho \cdot d\rho d\phi \\ 0 \end{pmatrix} \tag{5}$$

##### 4.1.1 Evaluation of each polarization intensity component

Of interest are the two polarization components defined according to the optical table horizontal and vertical axis which we align to the x and y directions of the dipole coordinate axis. Evaluation of the integrals in equation 5 gives

$$I_X(\rho_{max}, \Theta, \Phi) = A^2 \cdot \frac{n}{16\pi^2 \cdot f_{obj}^2 \cdot n_{bfp}} \cdot \bar{\mu}^\dagger \cdot H(\rho_{max}) \cdot \bar{\mu}, \tag{6}$$

$$I_Y(\rho_{max}, \Theta, \Phi) = A^2 \cdot \frac{n}{16\pi^2 \cdot f_{obj}^2 \cdot n_{bfp}} \cdot \bar{\mu}^\dagger \cdot V(\rho_{max}) \cdot \bar{\mu}, \tag{7}$$

where $H(\rho_{max})$ and $V(\rho_{max})$ are diagonal matrices with the diagonal elements $h_{ii}(\rho_{max})$ and $v_{ii}(\rho_{max})$. These elements are

$$\begin{aligned} h_{11}(\rho_{max}) = v_{22}(\rho_{max}) &= \tfrac{\pi}{4} \cdot \left(4 + \rho_{max}^2 + (\rho_{max}^2 - 4) \cdot \sqrt{1 - \rho_{max}^2}\right) \\ h_{22}(\rho_{max}) = v_{11}(\rho_{max}) &= \tfrac{\pi}{12} \cdot \left(4 - 3 \cdot \rho_{max}^2 + (\rho_{max}^2 - 4) \sqrt{1 - \rho_{max}^2}\right) \\ h_{33}(\rho_{max}) = v_{33}(\rho_{max}) &= \tfrac{\pi}{3} \cdot \left(2 - (\rho_{max}^2 + 2)\sqrt{1 - \rho_{max}^2}\right). \end{aligned} \tag{8}$$

##### 4.1.2 Evaluating total intensity

With these results we can calculate the total intensity which is the sum of both components

$$\begin{gathered} I_X(\rho_{max}, \Theta, \Phi) + I_Y(\rho_{max}, \Theta, \Phi) = \\ A^2 \cdot \tfrac{n}{16\pi^2 \cdot f_{obj}^2 \cdot n_{bfp}} \cdot \bar{\mu}^\dagger \cdot (H(\rho_{max}) + V(\rho_{max})) \cdot \bar{\mu}. \end{gathered} \tag{9}$$

Evaluating this expression we get the end result

$$I_{X+Y}(\rho_{max}, \Theta) = \\ A^2 \cdot \frac{n}{12\pi \cdot f_{obj}^2 \cdot n_{bfp}} \cdot \left(1 - \left(1 + \frac{\rho_{max}^2}{8}\left(1 + 3\cos(2\cdot\theta)\right)\right) \cdot \sqrt{1 - \rho_{max}^2}\right). \tag{10}$$

Note that the intensity only depends on the numerical aperture and the dipole polar angle $\Theta$.

#### 4.2 Simulating polarization distributions for different sliding models

The simulated polarization distributions in Extended Data Fig. 4d-f were generated by calculating polarization intensities according to the vector based dipole point spread function model (equations 6 and 7) while the dipole orientation $\bar{\mu}$ was sampled according to the different sliding models (uniform, linear or rotation coupled sliding) and experiment coordinate systems (polarization axis x aligned with average DNA direction or rotated 45° from it). Because of the isotropic excitation with unpolarized light in the camera experiments, the amplitude of the dipole moment is independent of $\bar{\mu}$ and $A$ was therefore set to 1. To account for shot noise, the total number of photons $n_{Tot}$ per time-step $t = 1$ ms in the simulations was sampled from a Poisson distribution (with a photon count rate of 1900 s$^{-1}$ according to the experimentally measured averages for LacI-R in the EMCCD experiments) while the number of photons $n_X$ and $n_Y$ in the two polarization channels was sampled from a binomial distribution, that is

$$n_X \sim B(n_{Tot}, \frac{I_X}{I_X + I_Y}), \tag{11}$$

$$n_Y = n_{Tot} - n_X. \tag{12}$$

In the uniform mode simulation, the protein has no preferred orientation during sliding and can rotate freely around the axis orthogonal to DNA and around the DNA itself. These rotations occur on a timescale much faster than the acquisition time of our camera-based experiments, which is implemented in the simulation by re-sampling $\bar{\mu}$ uniformly from the surface of the unit-sphere at every time-step. In the linear sliding mode the protein cannot rotate around any of its own, or the DNA axes. In the simulation the DNA was stretched along the x-axis and $\bar{\mu}$ is on average oriented in the direction of DNA, where slow wobbling of the fluorophore and DNA was simulated to obtain a $P_0$ distribution that approximately matches the experimentally LacI-R distribution in average and variance. Fluorophore wobbling was implemented by re-sampling $\bar{\mu}$ uniformly on a cap centered in the DNA direction that covers 45 % of the unit-sphere surface every 60 ms. DNA wobbling/anisotropy in stretching direction was implemented by re-sampling the DNA direction in the xy-plane from a normal distribution with a standard deviation of 15° every 600 ms. Every 600 ms a polarization signal was calculated according to equation 1 from the number of photons $n_X$ and $n_Y$ in the two polarization channels. 50 000 of such events is included in each simulated polarization distribution. The simulation of the rotational sliding mode was performed in the same way as the linear model but includes an additional fast rotation of the protein around the DNA axis with $D_r$ =130 000 rad$^2$s$^{-1}$ (see 4.3), effectively giving a rotation which is much faster than the acquisition time.

#### 4.3 Simulating fluorescence traces for rotating fluorophores

To simulate fluorescence intensity traces for flurorophores rotating diffusionaly, the dipole moment $\bar{\mu}$ was rotated around the DNA axis with an angle given by the mean squared angular displacement at every time-step $t$ of the simulation

$$\Delta\alpha = \pm\sqrt{2D_r t} \tag{13}$$

with equiprobable positive or negative rotation direction. The fluorescence intensity was then calculated from the vector-based point-spread function model (equation 10) at every time-step using the same excitation light a in the experiment, i.e.circularly polarized light in the xy-plane for SMCT-FCS. This was implemented by scaling the dipole orientation $\bar{\mu}$ by the length of its orthogonal projection onto the xy-plane i.e. setting $A = sin\Theta$.

#### 4.4 Pitch dependent autocorrelation model

In many experimental situations, the contribution from rotational Brownian motion on the autocorrelation function takes the form[8,9]

$$g(\tau) = C \cdot \left(1 + \sum_{i=1}^{5} c_i \cdot e^{-k_i \cdot \tau}\right) \tag{14}$$

where and $C$, $c_i$ and $k_i$ are constants dependent on the experimental setup and the rotating molecule. In the case for unconstrained 3D rotational diffusion of a spherical particle excited by linearly polarized light this relationship simplifies to

$$g(\tau) = C \cdot \left(1 + c \cdot e^{-6 \cdot D_r \cdot \tau}\right) \tag{15}$$

where $D_r$ is the rotational diffusion constant of the particle. In this study, the relationship between the autocorrelation function $g(\tau)$ and the pitch of the rotation coupled sliding was found by simulating fluorescence intensity traces emitted from dipoles rotating diffusionally around the x-axis (DNA). Brownian dynamics simulations of the rotation of the fluorophore were performed for both 1D (see 4.3) and 3D[10] rotational diffusion. After calculating the autocorrelation function from these intensity traces we found that the simulated autocorrelations could be described accurately with the model

$$g(\tau) = e^{-k \cdot D_r \cdot \tau} \tag{16}$$

where the model parameter $k = 4$ gave very good fits to the simulated data for rotation around one axis (Extended Data Fig. 7d), independent on the $D_r$ and numerical aperture used in the simulation. When the rotation of the particle is coupled to a translational movement, the relationship between the rotational diffusion constant $D_r$ and the translational diffusion constant $D$ can be found through the equations for mean-squared displacement $\langle \Delta x^2 \rangle$, and mean-squared angular displacement $\langle \Delta \alpha^2 \rangle$ respectively to be

$$D_r = \frac{D}{\left( \frac{\langle \Delta x^2 \rangle}{\langle \Delta \alpha^2 \rangle} \right)}. \tag{17}$$

The pitch of the rotation coupled sliding $p$, that is how many base pairs the protein moves per angle it rotates around the DNA, is defined as

$$p = \frac{\Delta x}{\Delta \alpha}. \tag{18}$$

After combining equations (16,17 and 18) a theoretical expression for the total autocorrelation function $g(\tau)$ for each sliding molecule can be written as

$$g(\tau) = h(\tau) \cdot e^{-\frac{kD\tau}{p^2}} + b \cdot f(\tau). \tag{19}$$

Where the background functions $h(\tau)$ and $f(\tau)$ contains all contributions to the total autocorrelation function that are independent of rotational sliding of the molecule and $b$ is the amplitude of the additive term of the background. In our experimental situation the autocorrelation function is normalized on a time interval $\tau_1 < \tau < \tau_E$, where the normalized autocorrelation function is defined as

$$g_N(\tau) = \frac{g(\tau) - g(\tau_E)}{\langle g(\tau_j) \rangle_n - g(\tau_E)} \tag{20}$$

where $g(\tau_E)$ is the value of the autocorrelation function at the end of the time interval and $\langle g(\tau_j) \rangle_n$is the average of the autocorrelation function at the first $n$ number of time lags on the time interval ($n = 2$ is used here). To minimize contributions from the background and any hypothetical contribution from DNA rotation, model fitting was done on the difference in autocorrelation between two constructs of LacI (LacI-MBP-R and LacI-R) sliding with different rates. If the two proteins translate along DNA with rotation coupled sliding with pitches $p_1$ and $p_2$, and translational diffusion constants $D_1$ and $D_2$, equations 19 and 20 give a theoretical expression for the difference in normalized autocorrelation. That is

$$\tilde{d}(\tau) = \frac{e^{-\frac{kD_1\tau}{p_1^2}} - e^{-\frac{kD_1\tau_E}{p_1^2}}}{\left\langle e^{-\frac{kD_1\tau_i}{p_1^2}} \right\rangle_n - e^{-\frac{kD_1\tau_E}{p_1^2}}} - \frac{e^{-\frac{kD_2\tau}{p_2^2}} - e^{-\frac{kD_2\tau_E}{p_2^2}}}{\left\langle e^{-\frac{kD_2\tau_i}{p_2^2}} \right\rangle_n - e^{-\frac{kD_2\tau_E}{p_2^2}}} \tag{21}$$

where it has been assumed that the contribuiton from rotation coupled slidinng to the total autocorrelation function is much higher than the contribution from the background (i.e. $e^{-\frac{kD\tau}{p^2}} >> h(\tau)$ and $e^{-\frac{kD\tau}{p^2}} >> b \cdot f(\tau)$). The theoretical expression for the difference in autocorrelation in equation 21 was then fitted to the experimentally measured differences $d(\hat{\tau_j})$ between the two proteins at all time lags $j$. Fitting was done by finding the least deviation weighted by the inverse of the standard deviations $\sigma_{i,j}$ of the differences in autocorrelation obtained from the single molecule data, that is by minimizing

$$s = \sum_{j=1}^{7} \frac{\|\tilde{d}(\tau_j) - \hat{d}(\tau_j)\|}{\sigma_j}. \tag{22}$$

Since LacI-R and LacI-MBP-R are both constructs of LacI, sliding with effective translational diffusion rates differing by only ~30% they are expected to have very similar effective pitches of rotation coupled sliding. We therefore simplify model fitting by setting $p = p_1 = p_2$ (giving only one parameter $p$ in the fitting). To account for errors associated with the estimation of translational diffusion constants, $D_1$ and $D_2$ are allowed to vary within their 95% confidence interval determined from the experimental data. Global minimization was performed by running the Levenberg-Marquardt algorithm 1000 times with randomly generated initial-values. Weights $\sigma_j$ as well as error estimates of the fitting parameters were obtained by bootstrapping, implemented by re-sampling the trajectories and re-calculating the difference in average autocorrelation $\hat{d}(\tau_j)$ at each time lag 2000 times and running the fitting algorithm on these samples. The described algorithm produced robust results by giving the same estimate of the pitch and the error associated with it when the algorithm was run repeatedly with different seeds for the random number generator.

The fitting method accurately returned the correct pitch estimates when the algorithm was tested on simulated data where the amplitude of the additative background was non-neglible. This was seen by setting the background function $f(\tau)$ to the experimental measured autocorrelations from stationary single molecules of LacI-MBP-R and LacI-R, and generating the theoretic autocorrelation functions according to equation 19 for different theoretical pitches. Since only the shape but not the amplitude of the background can be determined from the normalized autocorrelation of stationary molecules, simulations were performed at different amplitudes covering the interval which could have generated the experimental data (Extended Data Fig. 7 a,b). When autocorrelation functions were generated with $p_1 = p_2$ and with translational diffusion constants $D_1$ and $D_2$ set to the experimentally measured averages for LacI-MBP-R and LacI-R, the error in the pitch estimate was smaller than 7 bp for all relevant values off the background amplitude (Extended Data Fig. 7b), and smaller than 2 bp for estimated pitches close to experimentally observed estimated pitch.

#### 4.5 Stochastic simulations of LacI flipping and operator switching

To be able to determine the microscopic kinetics giving rise to our observed experimental data, we calculated the switching rate $s$ and flipping rate $f$ as a function of the model parameters $\Theta = (k_{on,\mu}, k_{off,\mu}, x_{hopp})$. This was done by simulating traces of operator switching and flipping. We then calculated the rates $s$ and $f$ by dividing the number of switching and flipping events by the total trace time, in the same way as we do for the experimental data. The algorithm for generating traces is described in the following pseudo code.

##### 4.5.1 Pseudo code for simulating traces

```
The time t = 0
Current operator = O_start
WHILE still running
    Sample the time to the next dissociation t_next from an exponential distribution with average
    1/(k_off,μ + k_off,macro)
    t = t + t_next
    n =random number sampled from uniform distribution between 0 and 1.
    IF n <k_off,μ/(k_off,μ + k_off,macro)
        The protein goes into the unspecific sliding/hopping mode.
        Given k_on and D_1^helix and the operator that the protein just dissociated from, sample the
        operator O_next, time t_bind and the number of hops for the next rebinding event from the
        precomputed lookup table (see 4.5.2). If the number of hops > 0, make the protein flip with
        a probability of 0.5.
        Sample the time t_dissociate for the next macroscopic dissociation from an exponential distri-
        bution with average 1/k_off,macro
        IF t_dissociate >t_bind
            Current operator = O_next
            t = t + t_bind
            Save the current operator, flip state, and t to the current trace.
        ELSE
            t = t + t_dissociate
            The protein dissociated macroscopically. End the current trace.
        ENDIF
    ELSE
        t = t + t_dissociate
        The protein dissociated macroscopically. End the current trace.
    ENDIF
ENDWHILE
```

To save computation time, the rebinding distributions (dependent on $k_{on}$ and $D_1^{helix}$ and $x_{hop}$) have been precomputed and saved in a look-up table before generation of the traces. The rebinding distributions are generated with the following algorithm.

##### 4.5.2 Pseudo code for computing the rebinding distribution

```
The time t = 0
Current position = position of O_start
WHILE still running

    IF the protein has met an operator

        r = k_on
        s = 2D_1^helix

    ELSE

        r = 0
        s = 2D_1^helix + k_hop

    ENDIF
    n =random number sampled from a uniform distribution between 0 and 1.
    t_next =random number sampled from an exponential distribution with average 1/(r + s)
    t = t + t_next
    IF n < r/(r + s)

        The protein binds the operator. Save t and the number of hops in the lookup table.

    ELSE

        The protein changes position
        n =random number sampled from a uniform distribution between 0 and 1.
        IF n < 2D_1^helix/(2D_1^helix + k_hop)
            Make helical sliding step, equidistributed either do the left or the right, and update the
            current position.
        ELSE
            Make a hop, equidistributed either to the left or the right. Sample the length of the hop
            from a normal distribution with standard deviation x_hop. Since the model assumes that
            the protein doesn't rotate in the hopping mode, the hop is rounded of to the closest 10
            bp to place the protein in the major groove an integer of DNA periods away from the
            last position. Update the current position and increase the hop counter.
        ENDIF

    ENDIF
ENDWHILE
```

The algorithm generated identical results (within numerical error) with 0.5 or 1 bp steps sizes in the helical sliding. To generate traces for $k_{on} = \infty$ and avoid instant rebinding to the operator, the protein is moved 1 bp away after each microscopical dissociation for this

$k_{on}$.

##### 4.5.3 Association rate increase caused by hopping

We also used simulations to investigate what effect the search process with combined helical sliding and hopping has on the rate of first encounter with the operator. Simulations were performed both for a model were the protein only slides helically with the diffusion rate $D_1^{helix}$, and for the hybrid model were the protein can also hop with the hop length and hop rate estimated from the experiments. To mimic biologically relevant conditions[3] the protein could dissociate macroscopically from the DNA with a rate off $k_{off,macro} = 1/0.003$ $s^{-1}$ in these simulations. When a protein search for a specific binding site on a piece of DNA that is much longer than the sliding distance, the sliding-hopping hybrid model reached the operator 100% more often, instead of dissociating from the DNA, compared to searching with helical sliding only. In the simulations this was implemented as a search on a 100 000 bp DNA where the protein starting position was sampled uniformly on the entire DNA. The 100% rate increase associated with the sliding-hopping hybrid search is explained by the substantially longer DNA distance that this model scans per non-specific association event, a property which compensates and outweighs the fact that the operator is sometimes bypassed during hopping (Supplementary Fig. 2).

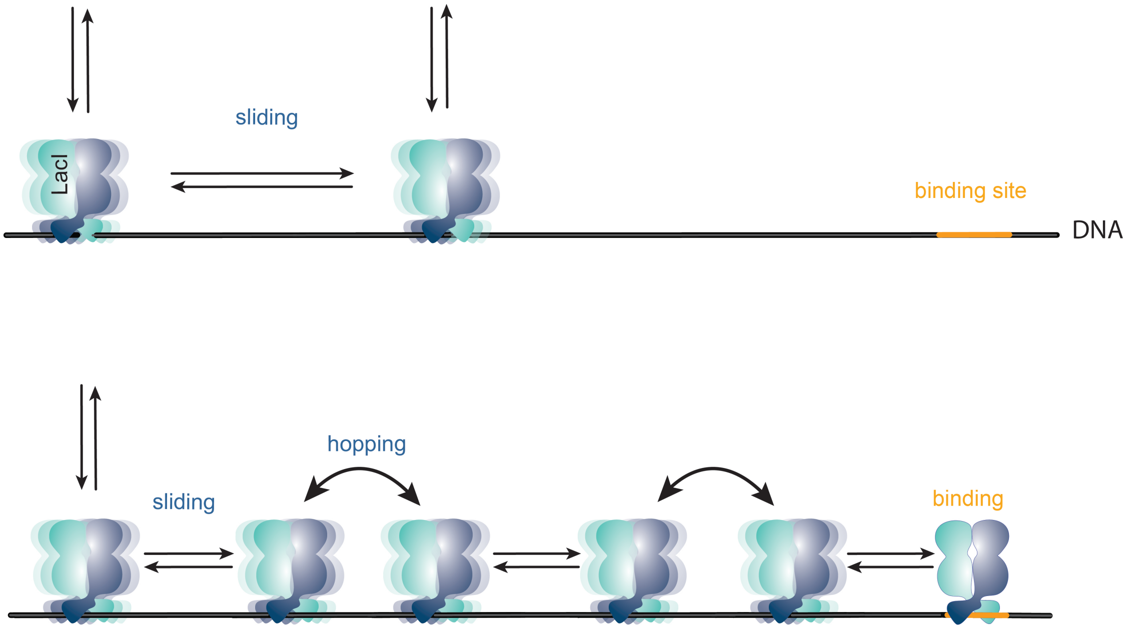


**Supplementary Figure 2.** Schematic illustration of operator search using helical sliding only compared to the sliding-hopping hybrid model.

##### 4.5.4 Probability densities of model parameters from experimental data

The probability for switching rates $s$ given traces $y$ is denoted $P_s(s|y)$ and can be generated from the experimental data by bootstraping over the experimental traces. Suppose that the values $s$ depends on hidden parameters $\Theta = (k_{on,\mu}, k_{off,\mu}, x_{hopp})$, meaning that the switching rate is a function of the model parameters, i.e. we have a function $s(\Theta)$. There now exists a probability distribution $P_s(\Theta|y)$, still associated to the variable $s$, which gives the probability of finding $\Theta$ given the traces $y$. The relation between $P_s(s|y)$ and $P_s(\Theta|y)$ is now given by

$$P_s(s|y) = \int_{\Theta \in s} P_s(\Theta|y) d\Theta \tag{23}$$

where $\Theta \in s$ means that we integrate over all $\Theta$ that give a specific $s$ value, the integral is thus over an isobar of $s(\Theta)$. We now assume that $P_s(\Theta|y) = P_s$ is constant over the parameter set of $\Theta$ values which gives one $s$ value. Restating this assumption we associate a constant probability $P_s$ for each isosurface of $s(\Theta)$ thus

$$P_s(s|y) = P_s \cdot \int_\Theta d\Theta = P_s \cdot V_{\Theta \in s} \tag{24}$$

The volume $V_{\Theta\in s}$ element can be reinterpreted as a function of $s$ , thus we can state that the volume $V_{\Theta\in s} = V(s)$. This gives us a relationship to calculate the sought probability density

$$P_s(\Theta|y) = \frac{P_s(s(\Theta)|y)}{V(s(\Theta))}. \tag{25}$$

In practice we chose to set $\Theta = (k_{off,\mu}, x_{hopp})$ while keeping $k_{on,\mu}$ fixed when estimating the mean and standard deviation of $x_{hop}$ from the experimental data, meaning that $V(s(\Theta))$ is a 2D area in $\Theta$-space instead of a 3D volume. This was repeated over the range of possible $k_{on,\mu}$ values to validate that the estimates are independent of this parameter, meaning that our results are equivalent to what would have been obtained if we solved the numerically more complex problem where $\Theta = (k_{on,\mu}, k_{off,\mu}, x_{hopp})$.
